# Structures of the heart specific SERCA2a Ca^2+^-ATPase

**DOI:** 10.1101/344911

**Authors:** Aljona Sitsel, Joren De Raeymaecker, Nikolaj Düring Drachmann, Rita Derua, Susanne Smaardijk, Jacob Lauwring Andersen, Ilse Vandecaetsbeek, Jialin Chen, Marc De Maeyer, Etienne Waelkens, Claus Olesen, Peter Vangheluwe, Poul Nissen

**Author notes:** Shared last and co-corresponding authors: Poul Nissen, Gustav Wieds Vej 10C, Aarhus University, 8000 Aarhus C, Denmark; Peter Vangheluwe, Campus Gasthuisberg, ON1, KU Leuven, Herestraat 49, box 802, 3000 Leuven, Belgium.

## Abstract

The isoform 2a of sarco/endoplasmic reticulum Ca^2+^-ATPase (SERCA2a) performs active reuptake of cytoplasmic Ca^2+^ and is a major regulator of cardiac muscle contractility. Dysfunction or dysregulation of SERCA2a is associated with heart failure, while restoring its function is considered as a therapeutic strategy to restore cardiac performance, but its structure was not yet determined. Based on native, active protein purified from pig ventricular muscle, we present the first crystal structures of SERCA2a that were determined in the CPA-stabilized and H^+^-occluded [H_2-3_]E2-AlF_4_ - (3.3 Å) form, arranged as parallel dimers, and the Ca^2+^-occluded [Ca_2_]E1-ATP (4.0 Å) form. We compare these new structures to similar forms of the skeletal muscle SERCA1a and address structural, functional and regulatory differences. We show that the isoform specific motifs of SERCA2a allow a distinct regulation by post-translational modifications and affect the dynamic behavior, which may explain specific properties and regulation.

## Introduction

The cardiac contractility is tightly regulated by the activity of the sarco/endoplasmic reticulum Ca^2+^ ATPase 2a (SERCA2a), which is responsible for the re-uptake of cytosolic Ca^2+^ into the sarcoplasmic reticulum (SR) of cardiomyocytes. SERCA2a enables cardiac muscle relaxation and determines how much Ca^2+^ can be released for contraction, which in turn controls contractile strength (Bers, 2002). In the heart, the activity of SERCA2a is regulated by several transmembrane (TM) micropeptides: Phospholamban (PLB) (MacLennan & Kranias, 2003), Sarcolipin (SLN) (Vangheluwe *et al*, 2005), Dwarf Open Reading Frame (DWORF) (Nelson *et al*, 2016) and Another Regulin (ALN) (Anderson *et al*, 2016). These small TM proteins regulate SERCA2a activity mainly via controlling the apparent Ca^2+^ affinity of the pump within a narrow physiological range thereby allowing a dynamic regulation of cardiac contractility. For instance, in resting conditions PLB inhibits SERCA2a, which is temporally relieved during β-adrenergic stimulation, and exerts strong inotropic and lusitropic effects (Arkin *et al*, 1994).

A reduced SERCA2a activity at least in part contributes to the progressive deterioration of cardiac contractility in heart failure (HF). Lower SERCA2a expression (Periasamy *et al*, 2008) and protein sumoylation (Kho *et al*, 2011), together with reduced levels of DWORF (Nelson *et al*, 2016) and PLB phosphorylation (Periasamy *et al*, 2008), negatively impact on the uptake of Ca^2+^ into the SR, impairing the contractile performance of the heart. Also, some familial forms of dilated cardiomyopathy are caused by mutations in PLB, further highlighting the central role of SERCA2a dysregulation in HF (Trieber *et al*, 2005; Haghighi *et al*, 2003; Kimura *et al*, 1998). Conversely, restoring SERCA2a activity by elevating its expression or by PLB interference rescues contractility in isolated cardiomyocytes, and in small and large animal models (MacLennan & Kranias, 2003). Therefore, adeno-associated viral gene delivery of SERCA2a was explored in clinical trials as a therapeutic strategy for HF (Jessup *et al*, 2011; Hulot *et al*, 2017), although inadequate SERCA2a gene delivery has been noted (Greenberg *et al*, 2016). Alternative to gene delivery, small molecule SERCA2a agonists have been reported (Ferrandi *et al*, 2013; Kaneko *et al*, 2017), but molecular details of their working mechanism are lacking.

SERCA2a (encoded by the *ATP2A2* gene) is expressed in cardiac, smooth and slow-twitch skeletal muscles, while SERCA1a (*ATP2A1*) is the major isoform in fast-twitch skeletal muscles. Overall, they are similar and transport two Ca^2+^ ions per ATPase cycle. Based on numerous SERCA1a crystal structures capturing most conformational states along the functional cycle, the SERCA Ca^2+^ transport mechanism is well described at the functional and structural level (Bublitz *et al*, 2013; Winther *et al*, 2013; Toyoshima *et al*, 2013). Three cytoplasmic domains (*i.e.* the nucleotide binding domain, N; phosphorylation domain, P; and actuator domain, A) regulate ATP binding, autophosphorylation and dephosphorylation, whereas a TM domain controls Ca^2+^ binding and translocation. During ion transport across the membrane, SERCA undergoes large conformational changes switching between Ca^2+^-bound E1 states and Ca^2+^-free E2 states (Moller *et al*, 2010; Post & Sen, 1965; Albers *et al*, 1963). In the E1 state, cytosolic Ca^2+^ enters via the Ca^2+^ entry gate and binds with high affinity at the TM domain. Mg^2+^- ATP binds to the N-domain, resulting in the phosphorylation of an aspartate residue in the P domain (Asp351, conserved in all P-type ATPases) and the formation of an occluded [Ca_2_]E1P intermediate. A spontaneous conformational transition to an E2P state, typically rate limiting for SERCA1a transport, opens the Ca^2+^ exit gate to the luminal compartment and allows Ca^2+^ release. Subsequently, negatively charged residues of the ion binding sites are partially protonated, which triggers occlusion and dephosphorylation, catalyzed by a conserved glutamate residue of the A domain (E183) via an E2-Pi intermediate (Olesen et al. 2004) and ending with the phosphate-released E2 state. The conversion of the protein to an E1 state opens the cytoplasmic Ca^2+^ entry gate, promoting the release of protons from the TM domain possibly via a C-terminal pathway (Bublitz *et al*, 2010) and allowing another cycle to begin (Winther *et al*, 2013; Toyoshima *et al*, 2013).

Compared to SERCA1a, heart muscle SERCA2a exhibits a lower maximal turnover rate and higher apparent Ca^2+^ affinity than SERCA1a (Dode *et al*, 2002, 2003). This implies that structural features somehow modify the kinetics of the Ca^2+^ transport mechanism. To study the SERCA2a specific properties, we purified porcine SERCA2a, which shares 99% sequence identity with the human SERCA2a and 84 % identity with rabbit and other mammalian SERCA1a. Subsequently, we determined structures of SERCA2a in a proton-occluded 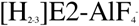 complex stabilized by the SERCA inhibitor cyclopiazonic acid (CPA) and a Ca^2+^-occluded E1 form using the non-hydrolyzable ATP analog AMPPCP. Furthermore, we compare SERCA1a and SERCA2a structures, and probe their regulation and dynamic behavior by molecular dynamics simulations, and address as well their functional differences using chimeric constructs.

## Experimental Procedures

### Isolation and purification of SERCA2a from pig heart

Pig hearts were collected from a slaughter house (Danish Crown in Horsens, Denmark) or obtained from the department of cardiovascular sciences (KU Leuven, Belgium), and were immediately rinsed with an ice-cold 0.9% (w/v) NaCl solution. All the following procedures were performed at 4°C. Ventricle tissue from the pig heart was homogenized with 10 mM NaHCO_3_ buffer (1:3 w/w) and centrifuged for 20 min at 12,200 × g. The supernatant was filtered through eight layers of gauze and centrifuged as above. The filtering step was then repeated and the supernatant was centrifuged at 140,000 × g for 45 min. The pellet was re-suspended in buffer (0.6 M KCl, 10 mM histidine, pH 7.0), magnetically stirred for 30 min, and then centrifuged again at 140,000 × g for 45 min. The final pellet containing crude SR microsomes was re-suspended in a stabilizing buffer (8 mM CaCl_2_, 50 mM MOPS-KOH, pH 7.0, 20% (vol/vol) added glycerol, 5 mM MgCl_2_) and flash-frozen in liquid nitrogen. A similar protocol preparing for purification by Reactive Red 120 beads was published earlier (Yao *et al*, 1998). The protein was solubilized in n-Dodecyl β-D-Maltopyranoside (DDM) at a ratio of 1:3 (w/w). Once solubilized, the samples were centrifuged for 45 min at 200,000 × g at 4°C. The supernatant was loaded onto Reactive beads (Green Separophore 4B-CL, Bio-world) and incubated for 1 h. Beads were washed with 5 column volumes of wash buffer (20% (vol/vol) glycerol, 20 mM MOPS-KOH pH 7.0, 1 mM CaCl_2_, 1 mg/ml C_12_E_8_; 0.35 mg/ml egg yolk phosphatidyl choline lipids (EYPC)). Finally, protein was eluted with 2 column volumes elution buffer (wash buffer with 50 mM NaCl and 4 mM AMPPCP). The protein concentration was measured using the Bradford protein assay (Bio-Rad). The presence of other SERCA isoforms was tested by Western Blot using the following antibodies: S2b AB (SERCA2b pAB serum, Badrilla); S2a AB (SERCA2a pAB serum, Badrilla) 1:100000; S2 AB (IID8) 1:2000.

### Activity measurements of purified SERCA2a and COS1 microsomes

For the activity measurements, the wash and elution buffers for the protein purification contained 0.25 mg/ml C_12_E_8_ and no EYPC was added. The Ca^2+^-dependent activity of the protein was measured using the Baginski assay probing inorganic phosphate released from ATPase activity (Baginski *et al*, 1967). Briefly, the reaction mixture containing 50 mM TES/TRIS, pH 6.9, 100 mM KCl, 7 mM MgCl_2_, 1 mM ethylene glycol-bis(β-aminoethyl ether)-N,N,N′,N′-tetraacetic acid (EGTA), 5 mM NaN_3_, 0.2 mM NaMoO_4_, 50 mM KNO_3_, and 150 ng (purified enzyme) or 5 *μ*g (COS1 microsomes) of the protein, as well as CaCl_2_ in different concentrations was initiated with 5 mM ATP (final concentration) and incubated at 37°C for 20 min. The reaction was stopped with the double reaction volume of ascorbic acid solution (170 mM ascorbic acid dissolved 0.5 N HCl and mixed with 4 mM NH_4_MoO_4_, solutions kept on ice). The samples treated with ascorbic acid solutions were incubated for 5 minutes on ice, and then triple reaction volume of colorimetric solution (210 mM Na_3_AsO_3_, 155 mM sodium citrate and 58 mM acetic acid) was added to stabilize the color. The absorbance was measured at 850 nm wavelength after an additional 5 minutes incubation at 37°C. The results were analyzed in Excel and Origin (OriginLab Corporation).

### Crystallization of SERCA2a

Purified SERCA2a was concentrated to 3.5 - 6 mg/ml and trace amounts of aggregated proteins pelleted by centrifugation at 100,000 × g for 10 min. Crystals were grown using the vapor-diffusion hanging-drop method, in which 1 µl of protein was mixed with 1 µl reservoir solution and equilibrated against 450 µl reservoir solution. For the CPA-stabilized E2-AlF_4_ - form, an egg yolk phosphatidylcholine (EYPC) to protein ratio of 1.7:1 (w/v) was used, and 200 *µ*M CPA, 80 mM KCl, 15 mM MgCl_2_, 2 mM EGTA and 10 mM NaF were added. The 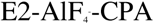 crystals were grown at 19° C for 4 days using reservoir solutions containing 14-20% PEG4000, 0-5% glycerol, 4-6% 2-Methyl-2,4-pentanediol and 100 mM Na^+^ malonate in the reservoir and then stored at 4°C. For the E1 conformation, the protein was treated with a final concentration of 10 mM CaCl_2_, 80 mM KCl and 3 mM MgCl_2_. [Ca_2_]E1-AMPPCP crystals appeared with reservoir solutions containing 21% PEG2000 monomethyl ether, 20% glycerol, 100 mM NaCl, 5% *tert*-BuOH and 2.4% Zwittergent-3-08 and grew within 7-10 days at 19°C. Crystals were fished with nylon loops. The E2-AlF_4_- CPA crystals were swiftly dipped in a mixture of 1 *µ*l reservoir solution and 1 *µ*l of 50% glycerol for cryoprotection. The mounted crystals were flash cooled and stored in liquid nitrogen.

### Data collection, processing and structural analysis

Diffraction data were collected at the Swiss Light Source (Villigen, Switzerland), at beam line PXI or PXII using a Pilatus 6M detector, with X-ray wavelengths at around 1 Å. Data were processed with XDS (Kabsch, 2010) to 3.3 Å maximum resolution in space group P2_1_ for the E2-AlF_4_- CPA form and to 4.0 Å in space group P2 for the [Ca_2_]E1-AMPPCP form. The structures were determined by molecular replacement using PHASER-MR (McCoy *et al*, 2007) and SERCA1a as a search model (PDB entries 3FGO and 3N8G) and revealing in both cases one SERCA2a molecule in the asymmetric unit. Model building was performed in COOT (Emsley & Cowtan, 2004) using electron density maps calculated using PHENIX (Adams *et al*, 2010). Refinement of the structural model was done in PHENIX (Adams *et al*, 2010). Both structures were built using the iMDff technique (Croll & Andersen, 2016; Focht *et al*, 2017). Structures were analysed with Molprobity, indicating for the E2-AlF_4_- CPA structure 98% residues in favored, 2% in further allowed, and 0 in non-favored region, with less than 1% rotamer outliers, and for the [Ca_2_]E1-AMPPCP structure 96% residues in favored, 4% in further allowed and 0 in non-favored regions of the Ramachandran plot, with no outlier rotamers. Figures were prepared and further model analysis performed with PyMol. The root mean square deviation for Cα atoms between the SERCA2a and the corresponding SERCA1a structures were determined by the ‘super’ command in PyMol.

### Molecular dynamics

Molecular Dynamics (MD) simulations were performed using the Gromacs package (version 4.5.3) (Hess *et al*, 2008). Each structure was placed in a dodecahedral box (x: 9.483 nm, y: 9.542 nm, z: 17.07 nm) and the system was solvated with TIP3P water molecules (Jorgensen *et al*, 1983). Lengths of x and y axis were fitted to the membrane dimensions. The energy of the system was minimized with 50,000 steps of steepest descent. Pre-equilibrated 1,2-Dioleoyl-sn-glycero-3-phosphocholine (DOPC) membrane structure and parameters were obtained from the Lipidbook website. The membrane system was energy minimized with 50,000 steps of steepest descent. Subsequently, all water molecules in the protein and membrane system were removed before merging them and inserting the protein into the membrane using g_membed tool in Gromacs. Next, the protein-membrane system was solvated with TIP3P and counter ions were added to neutralize the intrinsic negative charge of the SERCA2a structure. The energy of the protein-membrane system was minimized once more with 50,000 steps of steepest descent. The system was equilibrated using 500 ps of NVT ensemble with the V-rescale thermostat followed by 2 ns of NPT ensemble with the V-rescale thermostat and the Parrinello/Rahman barostat. Protein atoms and phosphorus atoms of lipid molecules were position restrained during equilibration with a force of 1000 kJ mol-1 nm-2. After equilibrium was reached, a full production MD without position restraint was performed for a time scale of 50 ns. In all simulations, an all atom AMBER99SB force field supplemented with AMBER-GAFF force field parameters for DOPC membrane (available on Lipidbook website) was employed.

### Plasmids, mutagenesis and cell culture

SERCA1a in pcDNA3.1 and SERCA2a in pMT2 were used for all expression experiments (Dode *et al*, 2003). Mutants were generated using the Quickchange site-directed mutagenesis kit (StrataGene) or the Q5 mutagenesis kit (New England Biolabs). A SERCA1a chimera with the luminal loop connecting transmembrane segments 7 and 8 (L7/8) of SERCA2a was previously described (Clausen *et al*, 2012). The reverse SERCA2a chimera with L7/8 of SERCA1a was generated using BBVC1 and BMGB1 restriction sites. Transfection into COS1 cells was performed with GeneJuice transfection reagent (Invitrogen). 72 h after transfection, the microsomal fraction was isolated by differential centrifugation as described previously (Maruyama & MacLennan, 1988).

### Mass Spectrometry

SERCA1a was purified according to a previously published protocol (Young *et al*, 1997). 10 *μ*g of purified SERCA2a and SERCA1a were reduced by 5 mM DTT and alkylated with 25 mM iodoacetamide, followed by precipitation (Wessel and Flügge 1984). The proteins were digested overnight using 0.5 *μ*g trypsin at 37°C in 200 mM ammonium bicarbonate, 5% acetonitrile, 0.01 % ProteaseMax (Promega). The resulting peptides were desalted with C18 ZipTip pipette tips (Millipore), and high-resolution LC-MS/MS on a 15 cm EASY-spray C18 column (Thermo Fisher Scientific) was performed using an Ultimate 3000 nano UPLC system interfaced with a Q Exactive hybrid quadrupole-orbitrap mass spectrometer. Peptides were identified by MASCOT (Matrix Science) using the SwissProt mammalian database via the Proteome Discoverer 2.2 software. Percolator was incorporated for peptide validation and ptmRS for PTM localization. Carbamidomethylation (C) was used as fixed modification, and acetylation (K), phosphorylation (STY) and ubiquitination (K) were each used in combination with oxidation (M) as variable modifications in the search parameter fields. Other search parameters included two allowed missed cleavages for trypsin digestion, peptide tolerance at 10 ppm and MS/MS tolerance at 20 mmu. Only peptides with a high identification confidence (PEP < 0.01) and a localization confidence of >99% were taken into account.

### Statistics

Results of the Ca^2+^ dependent ATPase measurements were fitted with the Hill function using Origin 8.0. One-way ANOVA with a Bonferroni post-hoc test was used for establishing significance.

## Results

### Purified SERCA2a from pig heart remains active

We first developed a one-step affinity chromatography protocol to purify native SERCA2a from pig left ventricular tissue, providing a yield of around 2.5 mg per 100 g of heart tissue (Figure 1A). In short, cardiac microsomes were solubilized with DDM, and SERCA2a was purified via Reactive Green affinity chromatography, which captures ATP-binding proteins. SERCA2a was subsequently eluted with AMPPCP. Immunoblotting experiments confirmed that the purified sample consists mainly of the SERCA2a isoform (Figure 1B). Only traces of the housekeeping SERCA2b isoform were found and no SERCA1a was detected (Figure 1B). No bands corresponding to PLB were detected on SDS-PAGE and immunoblotting (Figure 1B). The purified enzyme displayed robust, Ca^2+^-dependent ATPase activity, with a specific activity of 4.3 ± 0.7 *μ*moles/min/mg at 1 *μ*M free Ca^2+^, Hill coefficient n of 1.3 ± 0.1 and K_m_ value of 0.22 ± 0.01 *μ*M (Figure 1C). Thus, the purified SERCA2a from pig heart remains functional.

**Figure 1.**
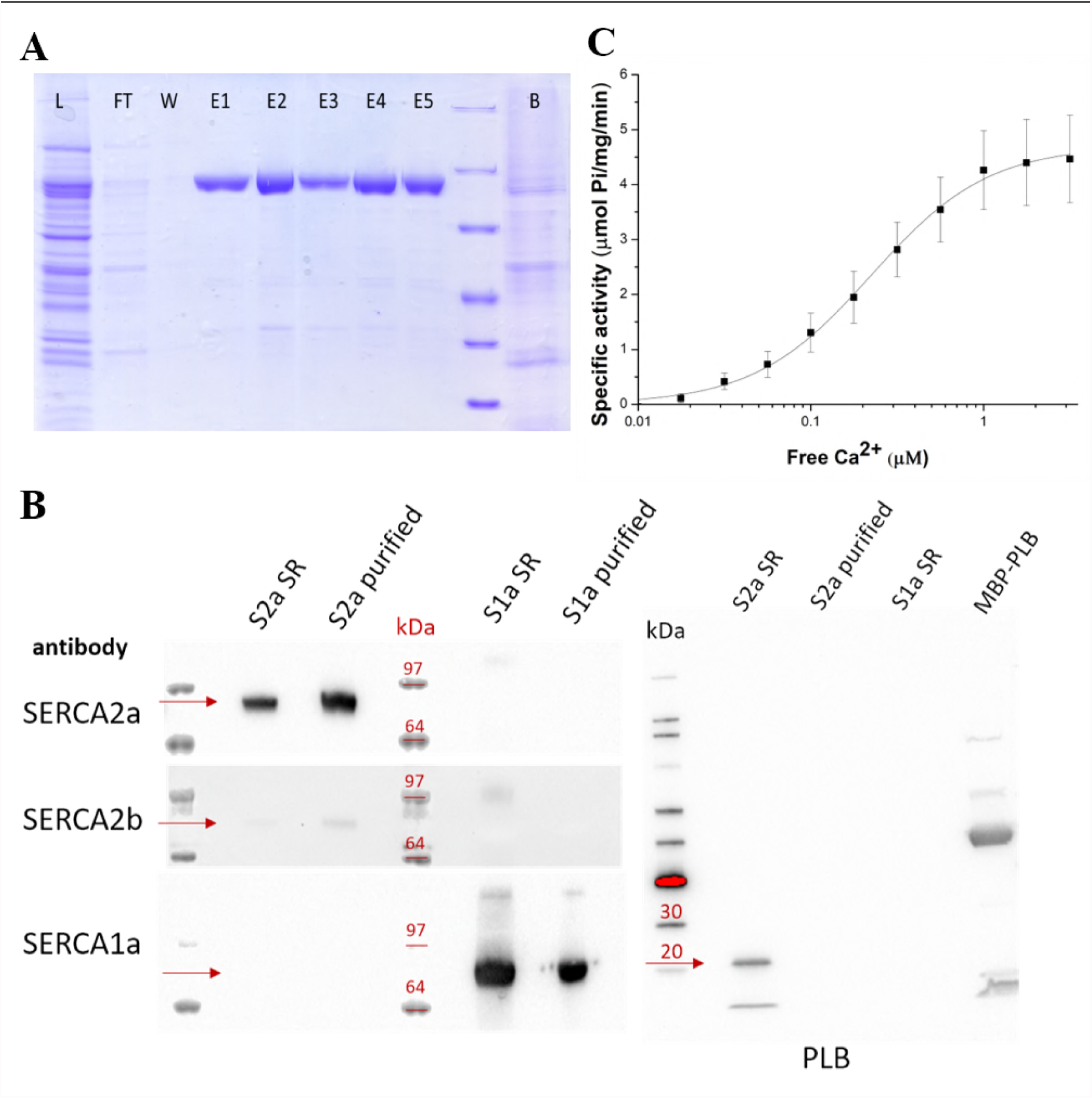
SERCA2a purification and activity. A. SDS-PAGE of fractions of the SERCA2a purification procedure (Load, total lysate; FT, flow-through from Reactive Green beads; Elution, elution fractions; Beads, beads after elution). B. Immunoblot with Sarcoplasmic Reticulum (SR) preparations of cardiac (S2a SR) and skeletal muscle (S1a SR), and purified SERCA2a (S2a purified) and SERCA1a (S1a purified). Detection was performed using antibodies against three SERCA isoforms 1a, 2a and 2b; or phospholamban (PLB). PLB appears as a monomeric (5 kDa) and pentameric band (25 kDa). A fusion construct of PLB with maltose binding protein (MBP) was used as a positive control for PLB detection. C. Ca^2+^-dependent ATPase activity of the purified SERCA2a. K_m_=0.22 ± 0.01 *µ*M, specific activity is 4.3 ± 0.7 *µ*moles/min/mg at 1 *µ*M free Ca^2+^, Hill coefficient n of 1.3.

### E2-AlF_4_- CPA and [Ca_2_]E1-AMPPCP structures of SERCA2a closely resemble SERCA1a

Structures of cardiac SERCA2a were determined in E2-AlF_4_- CPA and [Ca_2_]E1-AMPPCP forms at 3.3 Å and 4.0 Å resolution, respectively (Figure 2A, 2B and Table 1). In the [Ca_2_]E1-AMPPCP crystals, the TM regions of neighboring proteins interact in an antiparallel packing (*i.e.* not reflecting physiological contacts). However, the E2-AlF_4_-CPA crystal form is marked by an unusual parallel packing of SERCA2a molecules involving contact points between the A-domains and N-domains and between the A-domain and the P-domain domains, as well as N-domain and L7/8 (Supplementary Figure 1). No such parallel packing modes have been observed for SERCA1a crystal forms, and many of the involved residues differ for SERCA2a and SERCA1a and are conserved within one isoform. These SERCA2a specific regions may possibly play a role in the dimerization of SERCA2a in cardiomyocytes (Blackwell *et al*, 2016). Note that due to an amino acid deletion in SERCA2a at position 509, the SERCA2a residue numbering from position 509 is shifted by −1 as compared to SERCA1a.

**Figure 2.**
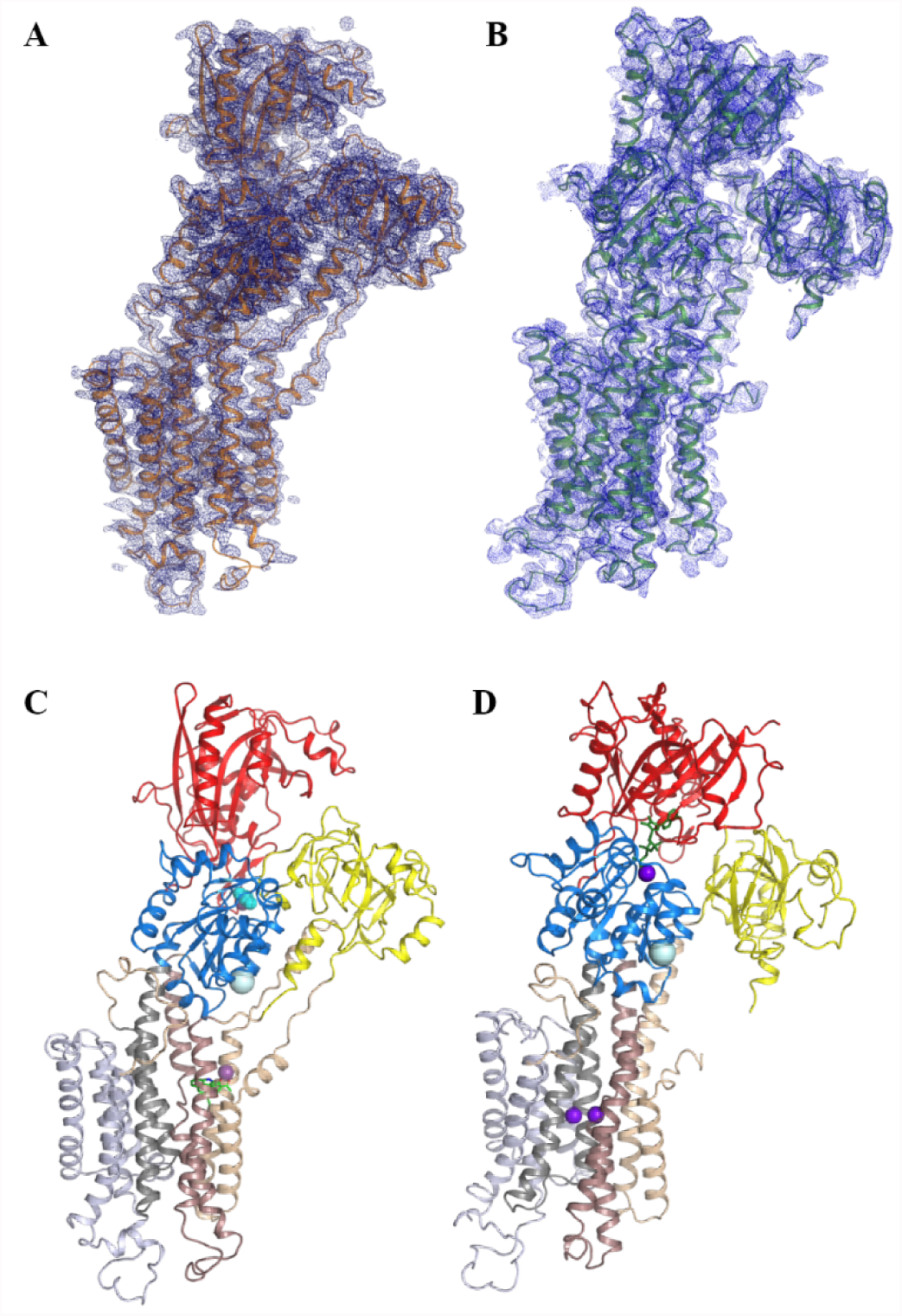
Crystal structures of cardiac SERCA2a in the 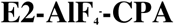 and [Ca_2_]E1-AMPPCP conformational states. Electron density maps of the SERCA2a E2-AlF_4_-CPA (A) and [Ca_2_]E1-AMPPCP (B) structures. Domain-colored SERCA2a in E2-AlF_4_-CPA (C) and [Ca_2_]E1-AMPPCP (D) states: A domain is colored in yellow, P in blue and N in red. M1-M2 are colored in wheat, M3-M4 in brown, M5-M6 in dark grey, and M7-M10 in light grey. Ca^2+^/Mg^2+^ ions are represented in purple spheres, AMPPCP in green sticks. The E2-AlF_4_-CPA structure depicts 992 amino acids of the 997 (no electron density was observed for the last five amino acids). The model of SERCA2a in the E1-Ca^2+^-AMPPCP conformational state contains 993 amino acids (Glu2-Gly994) of 997 total, with the exception of the residues Leu41-Leu49, Arg134-Lys135, Met239-Gln244, Ala424-Leu425, Ser504-Ser509, which were not build due to lack of electron density features.

**Table 1.**
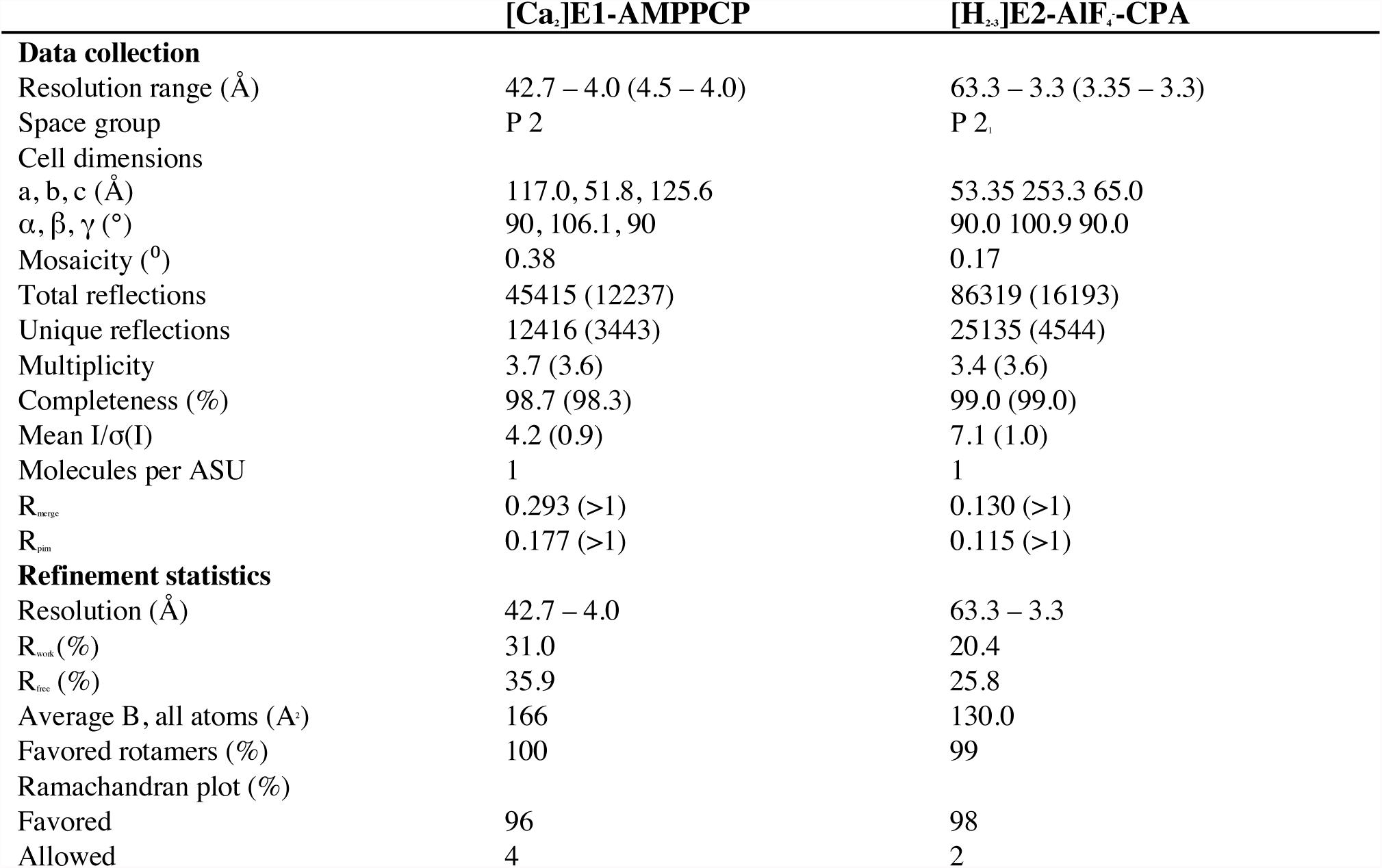
Data collection and refinement statistics for the SERCA2a structures in the Ca^2+^-E1-AMPPCP and 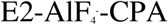 conformational states.

SERCA2a has the same overall domain organization as SERCA1a, including three cytoplasmic domains and 10 TM helices (Figure 2C, 2D). Indeed, the 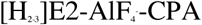 and [Ca_2_]E1-AMPPCP structures of SERCA2a are similar to the corresponding structures of SERCA1a (PDB 3FGO and 3N8G, respectively) (Figure 3A, 3B) with root mean square deviation (r.m.s.d.) for Cα atoms of 0.89 Å and 1.73 Å, respectively. Even the regions of highest sequence diversity, such as luminal loops L7/8 or L9/10 are similarly positioned in SERCA1a and SERCA2a. In both the E1 and E2 structures, the SERCA1a headpiece is tilted more towards the membrane as compared to SERCA2a. The [Ca_2_]E1-AMPPCP structure contains clear densities for the AMPPCP molecule as well as for bound Ca^2+^ and K+ ions. Ca^2+^ and AMPPCP binding in SERCA2a closely resembles the SERCA1a structure (Supplementary Figure 2). The CPA and MgF_x_ molecules as well as K^+^ ions are easily recognized in the 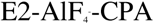 structure. However, AMPPCP is not observed despite being present at high concentrations in the crystallization conditions. Similar as in SERCA1a, CPA binds in a groove between helices M2, M3 and M4, coordinated by a Mg^2+^ ion and residues Gln56, Asp59 and Asn101, and blocks the proposed Ca^2+^ entry pathway (Laursen *et al*, 2009).

**Figure 3.**
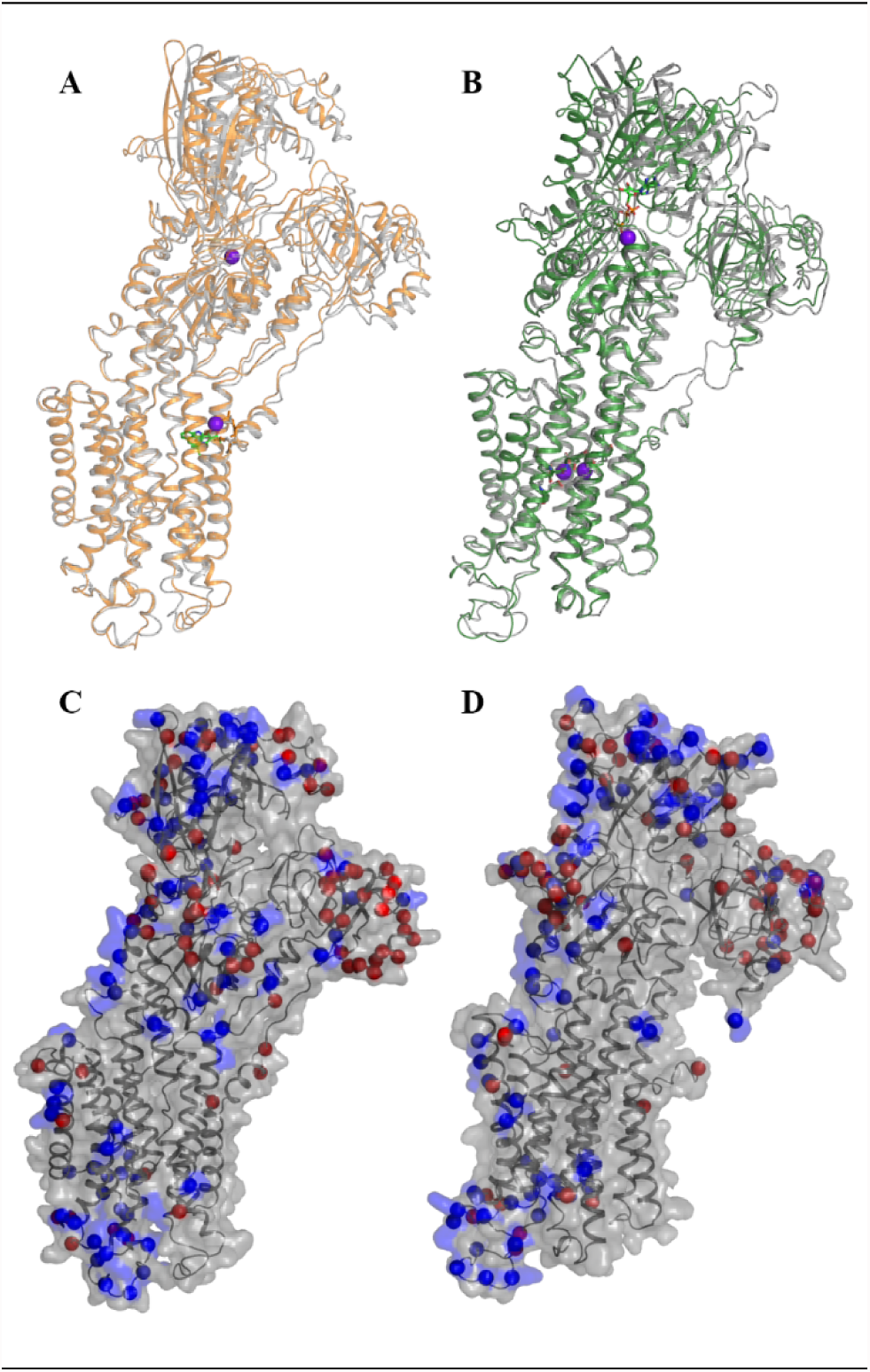
Comparison of the SERCA2a structures with the SERCA1a structures in the corresponding conformational states (PDB 3FGO and 3N8G) A. Alignment of the SERCA2a 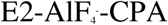 (orange) and SERCA1a (grey) structures. B. Alignment of the SERCA2a [Ca_2_]E1-AMPPCP (green) and SERCA1a (grey) structures C, D. Isoform specific residues are represented by both red and blue spheres mapped on the SERCA2a 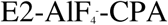 (C) and [Ca_2_]E1-AMPPCP (D) structures. Blue spheres depict residues that are identical in more than 90% of a selection of 83 vertebrate orthologues, while red spheres are less conserved.

### Isoform specific motifs of SERCA1a and SERCA2a

The catalytic core of the Ca^2+^ pump appears highly conserved (Supplementary Figure 3) indicating that the Ca^2+^ transport mechanism is preserved between SERCA1a and SERCA2a. Despite the 160 amino acid sequence differences between rabbit SERCA1a and pig SERCA2a, their structures do not readily reveal why the isoforms display distinct functional properties. As can be seen from Figures 3C and 3D, the isoform specific sequences are predominantly localized in exposed regions of the N-, P- and A-domains, near the membrane interface of the TM helices M7 and M10, and in the luminal loops L7/8 and L9/10. Based on an alignment of vertebrate SERCA1a and SERCA2a orthologues, many of these sequence differences were found to be highly conserved within one isoform (Supplementary Table 1), indicating that they may exert isoform-specific roles such as to serve as regulatory sites for specific protein or lipid interactions, or post-translational modifications (PTMs).

PLB, the major regulator of SERCA2a, is not present in the structure (Figure 1), but we examined the PLB binding site between M2, M4, M6 and M9 helices in the SERCA2a 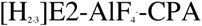 structure and compared it with the available SERCA1a-PLB and SERCA1a:sarcolipin structures (Akin *et al*, 2013; Winther *et al*, 2013; Toyoshima *et al*, 2013). The SERCA1a complexes adopt Ca^2+^-free E1 conformations, and the M2 therefore is shifted away by 9.4 Å in SERCA2a, but the relevant sidechains in the PLB binding groove are placed in a similar way (Leu802, Ala806, Phe809, Trp932, Leu939 which interact with Leu31 of PLB; and Gly801, Thr805 and Gln108, which interact with Asn34 of PLB), i.e. not pointing to major differences in the PLB acceptor site in the TM domain of SERCA2a and SERCA1a. We cannot exclude that specific interactions at the N-domain are Other SERCA2a regulators have been described (Vandecaetsbeek *et al*, 2009, 2011; Nelson *et al*, 2016; Anderson *et al*, 2015), but their interaction site are less well characterized.

### Isoform specific sites of post-translational modifications

SERCA isoforms are regulated by various PTMs, including sumoylation, phosphorylation, acetylation, glutathionylation, ubiquitination and nitration (Stammers *et al*, 2015). Interestingly, 26 out of the 160 isoform specific residues of SERCA2a (16%) are reported PTM sites supporting the view that many differences in the sequence may serve a regulatory function. Also, a higher number of unique PTMs has been reported for SERCA2a than SERCA1a (87 *versus* 29 reported PTMs). Mapping the previously reported PTMs onto SERCA2a and SERCA1a structures shows that the majority (79/87 in SERCA2a and 27/29 in SERCA1a) of PTMs are located in the cytoplasmic domains, accessible for enzyme interactions (Figure 4A, 4B) (Hornbeck *et al*, 2015).

**Figure 4.**
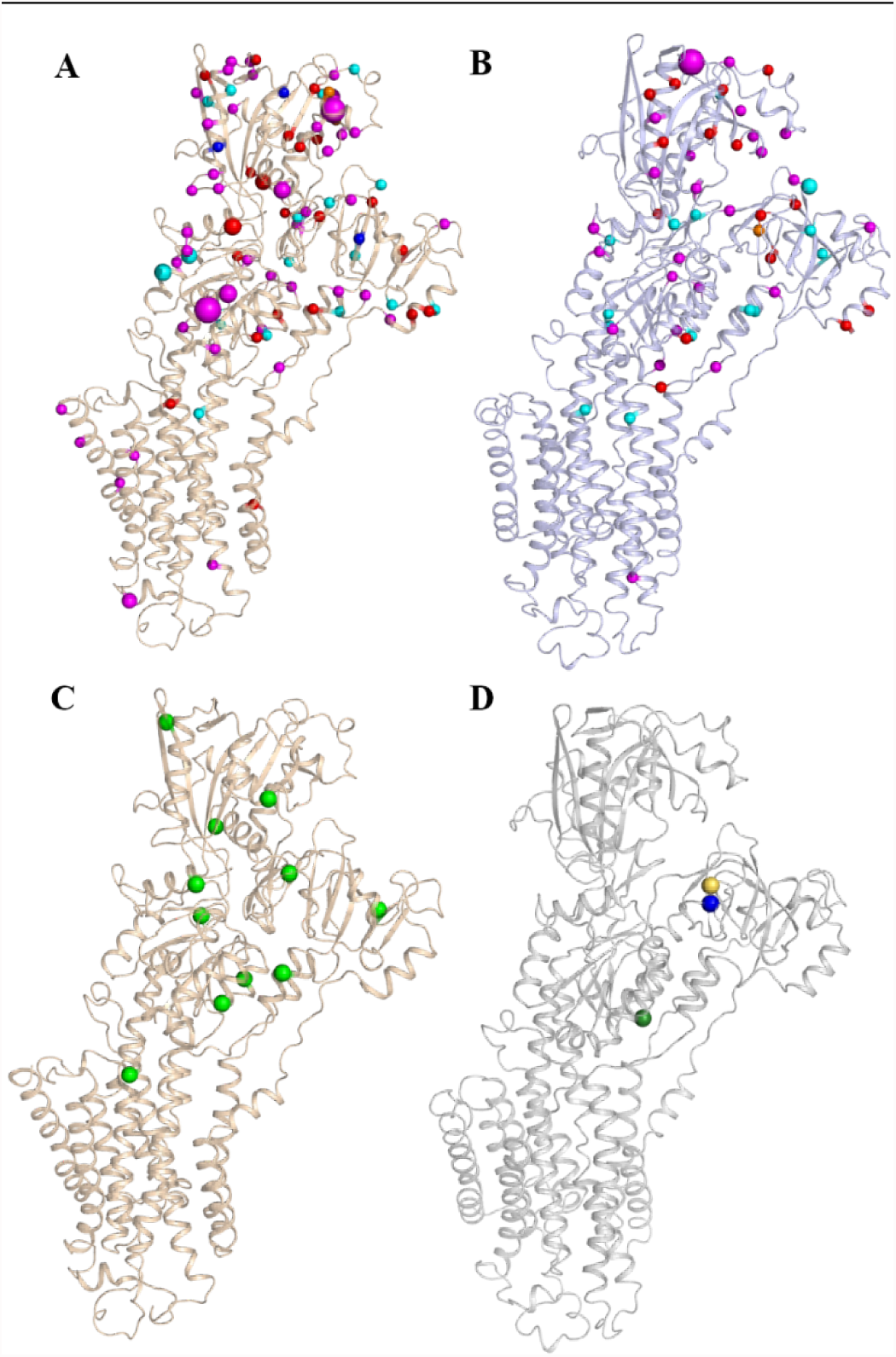
Post-translational modifications mapped on SERCA2a and SERCA1a structures. Post-translational modifications (PTMs) that were reported previously in human, mouse and rat (Hornbeck *et al*, 2015; Foster *et al*, 2013), as well as found in this study (in rabbit and pig) are shown in spheres for SERCA2a 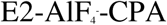 (A) and SERCA1a (PDB 3FGO) (B). Phosphorylation is shown in magenta spheres, ubiquitination in cyan, sumoylation in orange, acetylation in red and methylation in dark blue spheres. The size of the sphere correlates with the number of records obtained via a proteomic discovery-mode mass spectrometry approach (the smallest sized spheres correspond to PTMs that were identified one to five times, the medium sized sphere represents 6-25 hits and the largest sized spheres indicate >26 records). C. Newly identified acetylation sites found in purified SERCA2a in this study are shown in green spheres. D. Newly identified PTMs in purified SERCA1a by MS in this study: Lys204 (blue sphere) or Lys205 (yellow sphere) was found ubiquitinated, as well as Lys205 and Lys713 (green sphere) were found acetylated.

We experimentally determined which PTMs are present in our purified SERCA1a and SERCA2a samples. With an average coverage of 58.6% for SERCA1a and 60.3% for SERCA2a, MS analysis identified several new PTMs that have not been reported previously. Among these are multiple SERCA2a acetylation sites (Lys128, 205, 234, 352, 371, 451, 492, 628, 712, 727, 757) (Figure 4C, Table 2). In SERCA1a, we found two acetylated residues (Lys205 and 713) and one ubiquitination site (either Lys204 or 205) (Figure 4D, Table 2). However, despite the reported abundance of putative sites for post-translational control in SERCA2a, no additional electron densities were observed in the structures that may be ascribed to PTMs. The absence of PTMs in the electron density maps may be attributed to only low occupancy in the crystal, a high level of disorder, and/or a limited resolution of the structures, as also discussed for the palmitoylation of SERCA1a-SLN (Montigny *et al*, 2014). Additionally, unmodified SERCA2a may have been preferentially incorporated into the crystals during the crystallization process.

**Table 2.** Non-normalized (nmol P_i_/µg/min) and normalized (%) activity as well as the apparent Ca^2+^ affinity of the SERCA isoforms and chimeras. Mean values are given with standard deviation in parentheses. N=3 for each measurement.

|  | K<sub>m</sub> (μM) | V<sub>max</sub> (nmol P/μg/min) | V<sub>max</sub> (%) | n |
| --- | --- | --- | --- | --- |
| SERCA1a WT | 0.19 ± 0.04 | 0.51 ± 0.17 | 100.9 ± 2.1 | 1.5 ± 0.1 |
| SERCA1a-L7/8-2a | 0.16 ± 0.04 | 0.28 ± 0.08 | 80.1 ± 2.5 | 1.5 ± 0.2 |
| SERCA2a WT | 0.14 ± 0.02 | 0.18 ± 0.06 | 100.0 ± 2.8 | 1.5 ± 0.1 |
| SERCA2a-L7/8-1a | 0.10 ± 0.02 | 0.13 ± 0.07 | 71.6 ± 4.9 | 1.5 ± 0.1 |

### Isoform specific sequence differences alter molecular dynamics and functional properties

SERCA2a displays an intrinsically higher Ca^2+^ affinity and lower maximal turnover than SERCA1a (Dode *et al*, 2002, 2003), which should also be attributable to differences in the primary structure. To examine the functional role of the isoform specific sequence differences, we focused on the luminal loop between M7 and M8 helices (L7/8), which shows 15 substitutions on a total of 35 residues. It was shown before that a replacement of L7/8 in SERCA1a by the corresponding loop of SERCA2a (SERCA1aL7/8) alters the kinetic properties of the protein (Clausen *et al*, 2012). Here, we introduced the SERCA1a-specific L7/8 in SERCA2a (SERCA2a-L7/8-1a chimera) and the SERCA2a-specific L7/8 in SERCA1a (SERCA1a-L7/8-2a chimera) and compared the functional properties of these chimeras with SERCA1a and SERCA2a WT in a COS overexpression system. Expression levels for all constructs were similar (Supplementary Figure 4). We did not observe significant changes in the K_m_ values of the chimera *versus* their corresponding WT protein, but the substitutions of L7/8 in each isoform significantly reduces the Vmax by 20-30% (Table 2). While the L7/8 sequence differences alone do not explain the higher V_max_ and K_m_ of SERCA1a as compared to SERCA2a, the results show that the sequence of L7/8 seems to be only optimal in its own protein core. This suggests that the distinct functional properties of SERCA2a and SERCA1a relate to a complex interplay between sequence differences at various positions of the protein.

In the absence of clear structural differences between SERCA2a and SERCA1a, we considered that the distinct functional properties of both isoforms may rather relate to different dynamics. To explore this possibility, we compared 50 ns MD simulations of SERCA2a 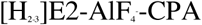 and its corresponding SERCA1a structure inserted in phosphatidylcholine membranes. The results suggest that isoform specific residues alter intramolecular salt bridge and hydrogen bond interactions that affect the protein dynamics (Supplementary Table 3). Indeed, a significant percentage of salt bridges or hydrogen bonds that are unique to each isoform encompass isoform specific residues: 42.9% in SERCA1a and 28.1% in SERCA2a for salt bridges, and 28.9% of SERCA1a and 39.5% of SERCA2a for hydrogen bonds. Hence, isoform specific residues are overrepresented in distinct networks. With respect to L7/8, we found that residues Val283, His284, and Gly285 of L3/4 in SERCA2a dynamically interact with Lys876 and Asn879 of L7/8 (Figure 5A). These interactions differ from SERCA1a, where Arg290 (L3/4) interacts with Thr877 and Glu878 (L7/8), and Ser287 (L3/4) interacts with the Gln875 and Glu878 (L7/8). Furthermore, residues Glu83 (L1/2), Glu892 (L7/8) and Glu895 (L7/8) of SERCA1a interact with SERCA1a-specific residues of L9/10 (Lys958, Lys958 and Lys960, respectively) (Figure 5B). In both isoforms, Gln965 (Gln966 in SERCA1a) of L9/10 is involved in hydrogen bond interactions with isoform specific residues of L7/8 (Glu895 and Asp861 in SERCA1a and Tyr894 and Gly860 in SERCA2a)with different interaction times (72% and 26%, respectively). (Figure 5A, 5B). Thus, the isoform specific dynamic behavior of L7/8 depends on interactions with neighboring loops L3/4 and L9/10, which may explain why replacing L7/8 in both the SERCA1a-L7/8-2a and SERCA2a-L7/8-1a chimeras is associated with a loss of function.

**Figure 5.**
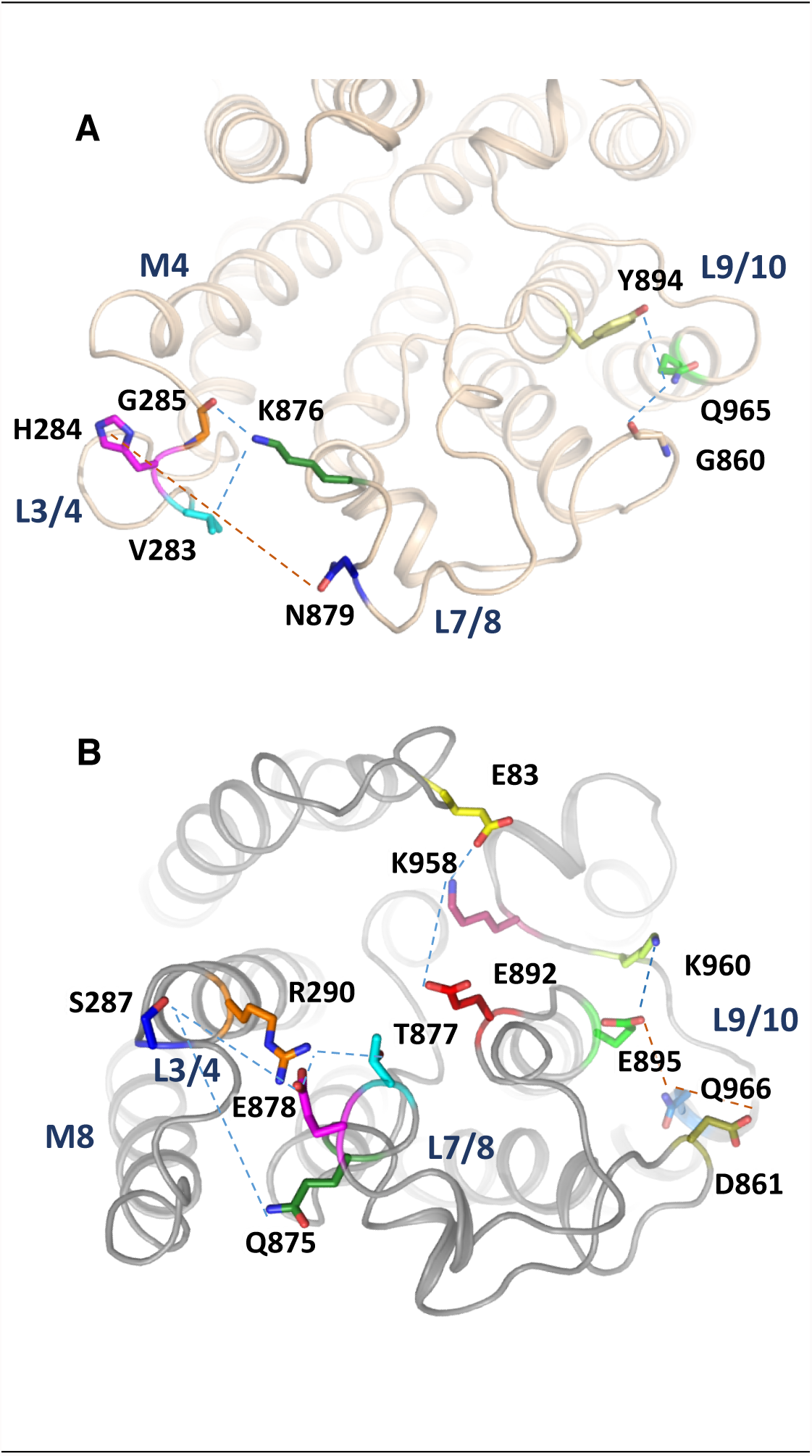
Residues of the luminal loops involved in isoform specific interactions according to molecular dynamics simulation. A. SERCA2a in the 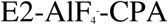 state depicting SERCA2a-specific residues involved in isoform unique interactions during MD simulations. B. SERCA1a (PDB 3FGO) depicting SERCA1a-specific residues involved in isoform unique interactions during MD simulations. The two panels are shown from the luminal side on the membrane, but in slightly different orientations for clarity on involved residues.

## Discussion

In this study, we developed a high yield purification protocol of native SERCA2a from pig heart and determined the first crystal structures representing two conformational state. We combined structural information, MD simulations, mass spectrometry, and a biochemical analysis to compare it to the muscle specific Ca^2+^-ATPases, SERCA1a.

### Role of SERCA2a and SERCA1a specific sequences

The good resemblance between SERCA2a and SERCA1a structures is consistent with a highly conserved Ca^2+^ transport mechanism in the two isoforms. However, we show that the sequence variations in SERCA2a and SERCA1a are functionally relevant, alter the dynamics, and allow isoform specific regulatory control. The conserved structure indicates that it may be possible to study the functional role of the isoform sequences by generating chimera proteins of SERCA1a and SERCA2a. However, we notice that the functional effect of isoform specific regions depends on the structural context, which is also isoform specific. We further depict that many sites for PTMs are located in exposed regions of the pumps and are isoform specific, indicating that these unique sequences support regulatory control. It is well established that SERCA2a is regulated by an increasing number of PTMs, and that this regulatory control may be disturbed in pathological conditions such as HF (Kho *et al*, 2011). For instance, the progressive accumulation of SERCA2a tyrosine nitration has been described during aging (Viner *et al*, 1999) and SERCA2a nitration levels are clearly correlated with HF (Lokuta *et al*, 2005). Moreover, nitration of SERCA2a-containing microsomes has a strong negative effect on Ca^2+^ uptake (Lokuta *et al*, 2005), and sumoylation regulates SERCA2a activity and stability and is decreased in HF, while restoration of SERCA2a sumoylation provides cardioprotection (Kho *et al*, 2011). An increased SERCA2a acetylation is associated with higher activity and altered intracellular Ca^2+^ dynamics influencing performance of cardiomyocytes (Meraviglia *et al*, 2018). The growing number of PTMs represents a major challenge to decipher the functional impact of each individual modification. A future assessment of the SERCA2a PTM profile of SERCA2a isolated from healthy and diseased hearts will allow us to establish a fingerprint of different SERCA2a PTMs in disease and pinpoint their effect on local and global dynamics of SERCA2a.

One of the most varying regions between SERCA1a and SERCA2a at the luminal side of the membrane is L7/8, which appears to be an important regulatory site. SERCA isoforms are regulated by a disulfide bridge formed between two conserved Cys residues on L7/8 (Daiho *et al*, 2001; Ushioda *et al*, 2016). In SERCA1a, the disulfide bridge inactivates the pump activity, while in SERCA2b, the reversible formation of a disulfide bridge between C875 and C887 serves as a luminal redox sensor that is controlled by an ER disulfide reductase (Ushioda *et al*, 2016). Furthermore, binding of calumenin to L7/8 leads to a reduction in the apparent Ca^2+^ affinity of the ATPase (Sahoo *et al*, 2009). In addition, only in SERCA2 isoforms L7/8 serves as an acceptor site for the luminal extension of the SERCA2b C-terminus, indicating that L7/8 is important for the isoform specific properties. The functional analysis of this study shows that a simple replacement of L7/8 reduces the maximal turnover rate in both the SERCA1a and SERCA2a backgrounds. According to previous kinetic measurements, L7/8 of SERCA2a lowers the Ca^2+^ dissociation rate towards the cytosol, and the rate of the E2 to E1 conversion is also reduced (Clausen *et al*, 2012). Together this suggests that L7/8 is adapted to the local environment in each isoform, which is confirmed by the MD simulations that highlight dynamic interactions between L3/4 and the isoform specific loops L7/8 and L9/10.

### Mapping Darier disease mutations on the SERCA2a structure

Several mutations in the *ATP2A2* gene are associated with Darier’s disease (DD), an autosomal dominant skin disorder, which is characterized by keratosis, skin and nail defects (Sakuntabhai *et al*, 1999b). Here, we used the SERCA2a structures to map all known DD mutations of various disease impact (Stenson *et al*, 2017). The map demonstrates that DD mutations are scattered all over the protein, without specific clusters that correlate with disease severity (Figure 6). DD mutations affect SERCA2a/b activity by a reduced protein stability and/or impaired functionality, which point to haploinsufficiency explaining the dominant inheritance pattern. Although DD mutations affect all three SERCA2 splice variants (SERCA2a-c), clinical manifestations only appear in the skin, where the ubiquitous SERCA2b isoform is expressed, while cardiac functionality is surprisingly well preserved (Tavadia *et al*, 2001; Mayosi *et al*, 2006).

**Figure 6.**
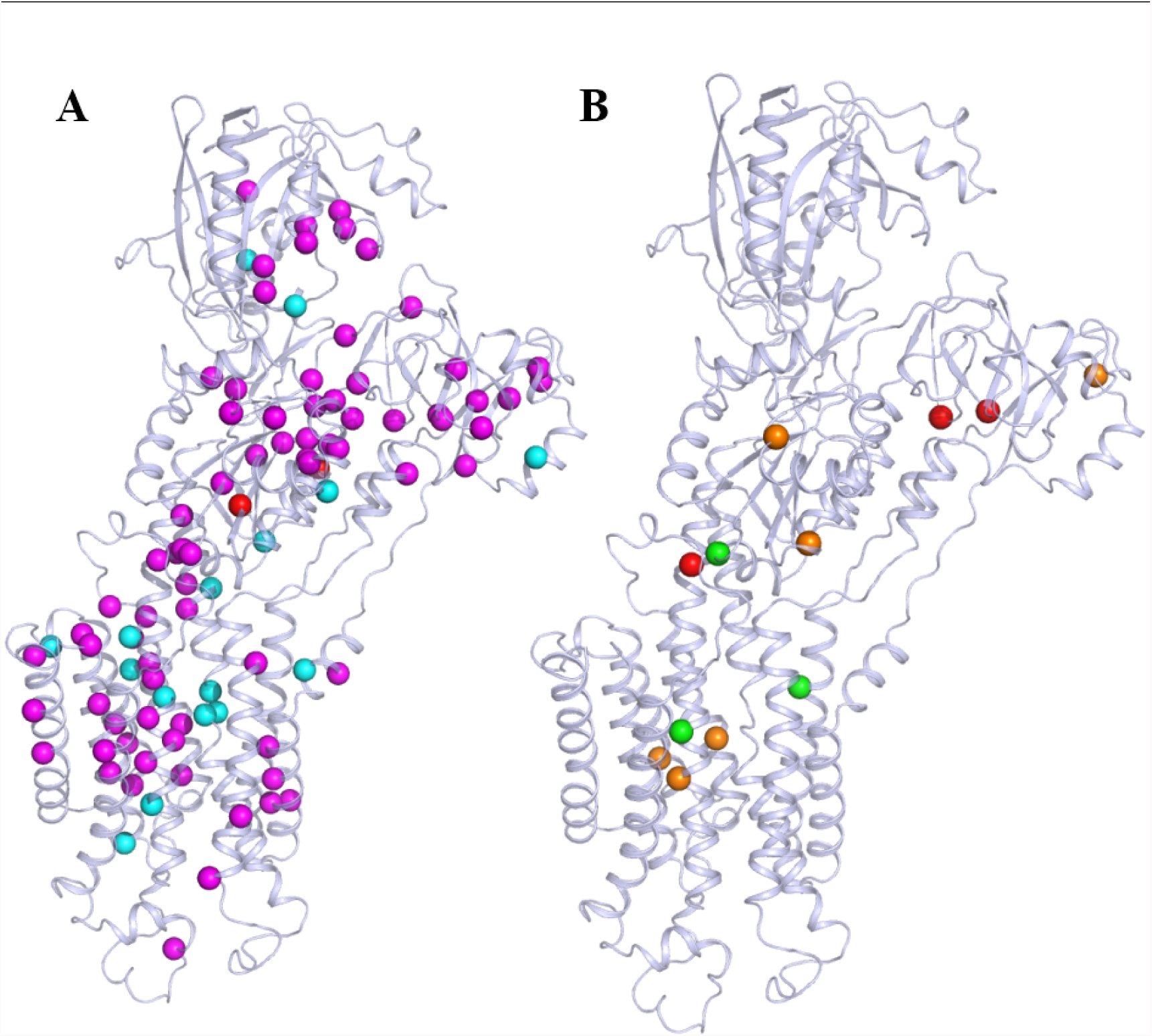
Darier-disease mutations. A. Darier disease (DD)-associated mutations shown in magenta spheres are missense/nonsense mutations, in cyan are most likely DD mutations and in red are associated with acrokeratosis verruciformis/hypertension. B. The review of Nellen *et al.* correlates DD mutations with the severity of the phenotype (Nellen *et al*, 2017). Mutations leading to a mild phenotype are represented by green spheres, moderate by orange and severe by red. Some of the mutations with the described severe (Pro160, Gly211, Gln691, Arg750, His943) and moderate phenotypes (His943, Asn795, Gly769, Gln691, Ala672, Arg131) are associated with lower expression and activity levels of the protein (e.g. Arg750, His943 and Arg131) (Sorensen & Andersen, 2000; Ruiz-Perez *et al*, 1999; Miyauchi *et al*, 2006; Sakuntabhai *et al*, 1999a; Ringpfeil *et al*, 2001; Chao *et al*, 2002). Also, Gly211 is related with the slower turnover rate of the protein and Gly769 with a higher affinity and Ca^2+^ transport (Ruiz-Perez *et al*, 1999; Miyauchi *et al*, 2006; Ren *et al*, 2006).

## Conclusion

The first structures of the SERCA2a isoform presented here represent important steps to facilitate further structural and functional studies of SERCA2a in normal and diseased context. SERCA2a is a favorable and recognized target for HF therapy, and the study also opens the door for the future crystallization of SERCA2a complexes with small molecules or TM regulators like PLB, which will aid drug discovery efforts. The SERCA2a and SERCA1a structures are very similar, which indicates a conserved mechanism of Ca^2+^ transport, but the isoform specific sequence differences introduce unique sites of regulation and may alter the molecular dynamics and kinetic behavior of the pump in different tissues. These insights may be of interest for drug discovery efforts to develop SERCA2a specific modulators, which will also be facilitated by the availability of the SERCA2a structural information.

## Acknowledgements

We would like to thank Maike Bublitz and Jesper Lykkegaard Karlsen for their help with data processing and iMDff experiments. We are grateful to Marleen Schuermans, Ingrid Puusta, Boyin Liu, Tugce Arslan, Anne Lindeman, Anna Marie Nielsen, Lotte Thue Pedersen and Tetyana Klymchuk for their excellent technical assistance. Also, we would like to thank Howard Young for the initial input to establish a purification protocol. We would like to thank the Swiss Light Source (Paul Scherrer Institute) for providing data-collection facilities, and the EMBL beamlines at ESRF and DESY for crystal screening. This work was funded by the Flanders Research Foundation FWO (G044212N and G0B1115N), and the Inter-University Attraction Poles program (P7/13) assigned to PV. The work was supported by an ERC grant (BIOMEMOS) to PN. The DANDRITE center is funded by the Lundbeck Foundation (grant no. R248-2016-2518). AS was supported by the IWT doctoral scholarship provided by Agentschap Innoveren & Ondernemen (VLAIO).

## Supplementary

**Supplementary Figure 1.**
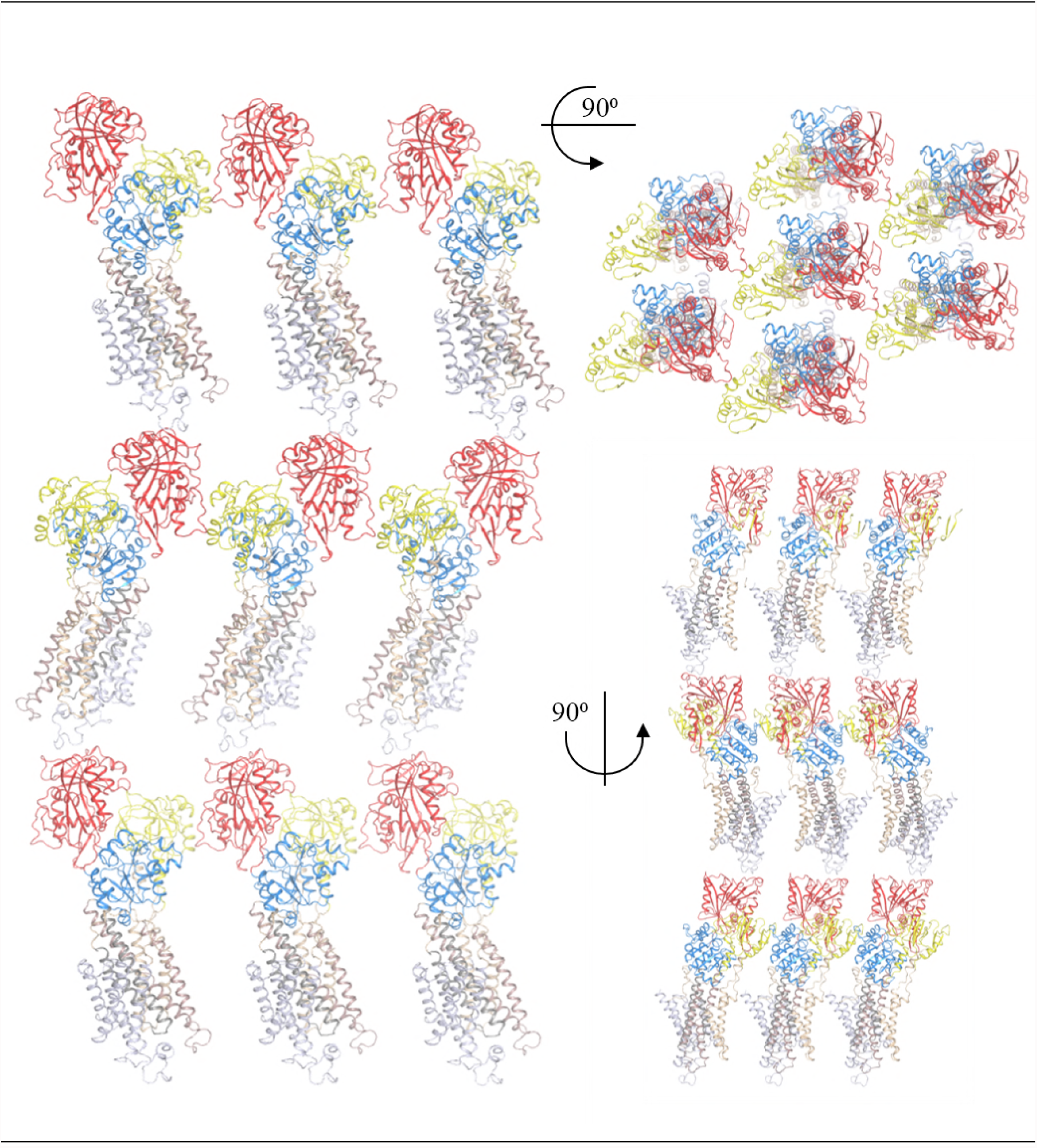
Packing of SERCA2a 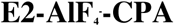 crystals. Parallel packing of the SERCA2a structure in the 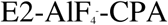 conformational state. SERCA2a domains are colored the same as in Figure 2B and 3B. SERCA2a forms contact points between the A-domains and N-domains (Met1-Asn3, His15 to Glu458-Lys460) and between the A-domain (Arg134, Arg139, Lys141) and the P-domain (Glu659, Glu667, Ser663), as well as N- (Thr506-Ser509, Asp567-Arg571) domain and L7/8 (Asp859-Arg863, Lys876-Gly884).

**Supplementary Figure 2.**
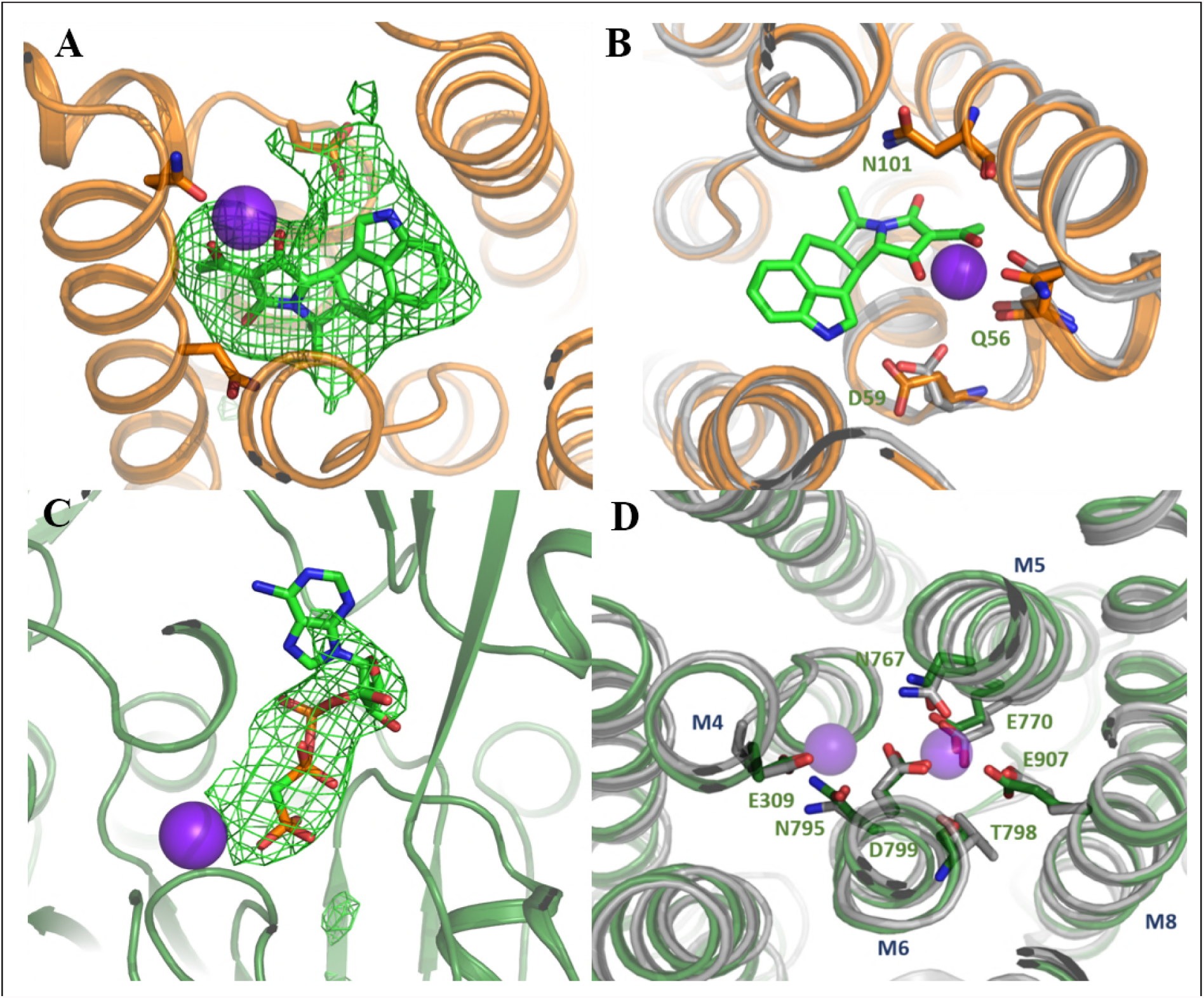
Coordination of ions and molecules in the SERCA2a structures. Ca^2+^ ions are shown in purple spheres. Omit maps for the CPA (A) and the AMPPCP (C) molecules contoured at 3σ. B. CPA coordination in 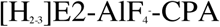 state of SERCA, SERCA1a is depicted in grey, SERCA2a in orange. D. coordination of Ca^2+^ ions in the [Ca2]E1-AMPPCP state. SERCA1a is shown in grey, SERCA2a in green. Residues are marked according to the SERCA2a numbering.

**Supplementary Figure 3.**
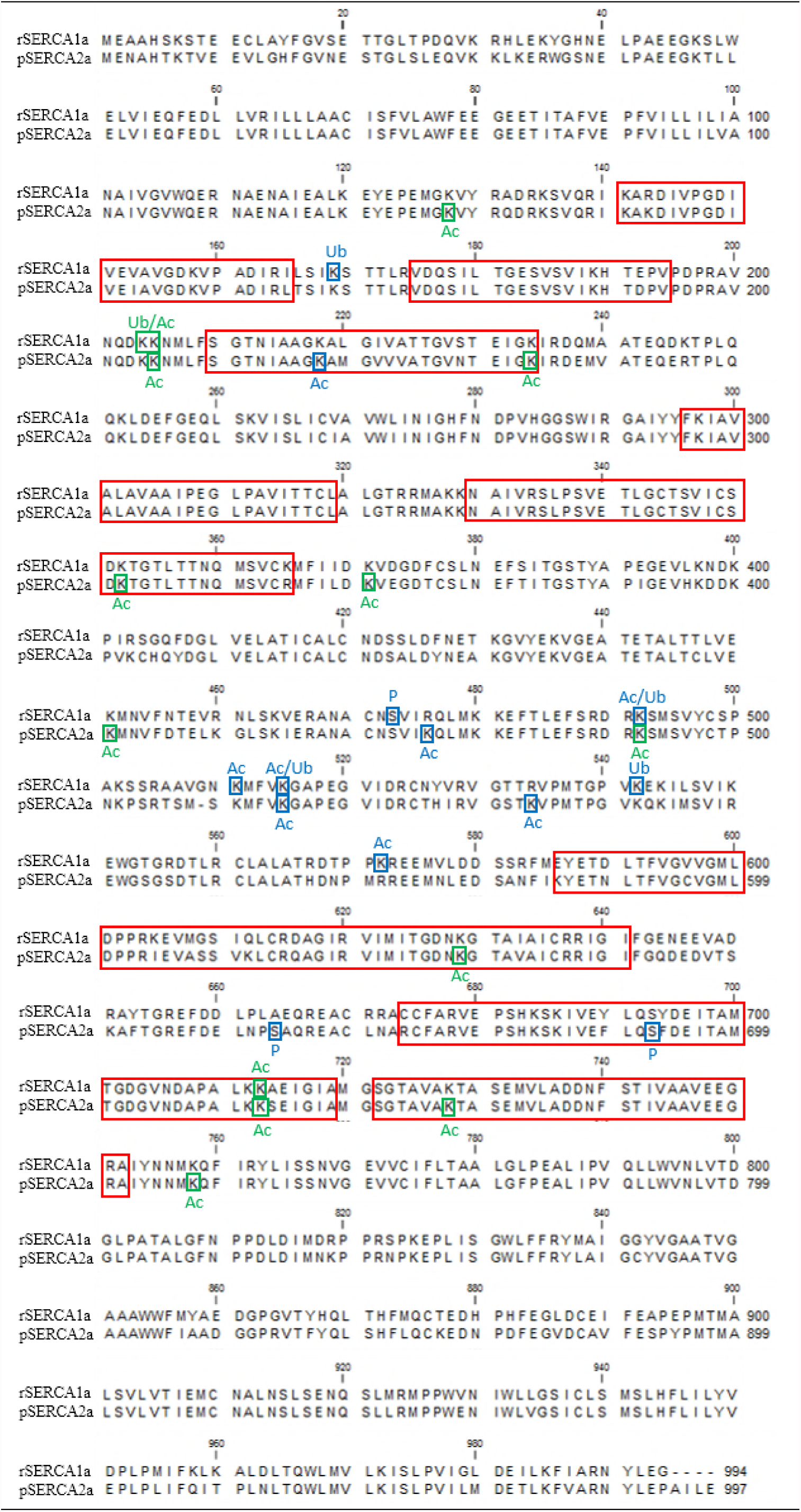
Sequence alignment of SERCA1a and SERCA2a. Alignment of rabbit SERCA1a and pig SERCA2a protein sequences indicating the eight most conserved regions in P-type ATPases (red squares) (Axelsen & Palmgren, 1998). The post-translational modifications found by MS in this study are marked by blue (reported before) and green squares (identified in this study).

**Supplementary Figure 4.**
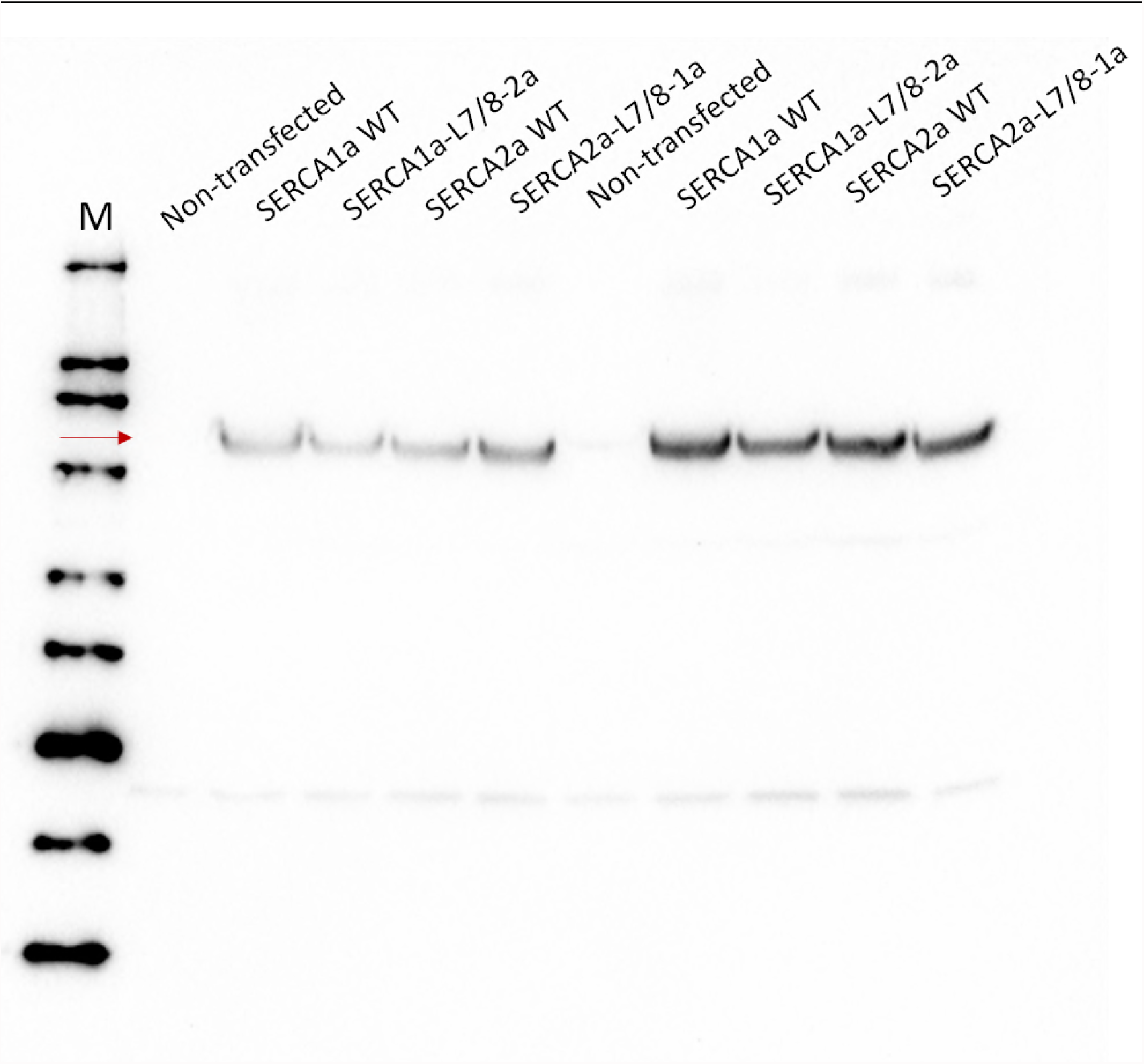
Expression levels of SERCA1a and SERCA2a constructs. Immunoblot with a non-isoform specific SERCA TRY2 antibody (Mountian *et al*, 2001) depicting similar protein levels of the WT and chimera constructs of SERCA2a and SERCA1a following transient expression in COS cells.

**Supplementary Table 1.** Conservation of the SERCA1a and SERCA2a specific residues based on 68 and 83 vertebrate sequences, respectively. For SERCA1a, all of the distinct residues in rabbit SERCA1a and pig SERCA2a are at least 39% conserved and 79% are at least 70% conserved, 12.5%. For SERCA2a, all of the distinct residues are at least 43% conserved, while 89% are at least 70% conserved. Rabbit S 1a RN – rabbit SERCA1a residue name, rabbit S1a RN – SERCA1a residue number pig S2a RN – pig SERCA2a residue name pig S2a RN – SERCA2a residue number.

| **Rabbit S1a RN** | **Rabbit S1a RN** | **Conservation (%)** | **Substitutions** | **Pig S2a RN** | **Pig S2a RN** | **Conservation (%)** | **Substitutions** |
| --- | --- | --- | --- | --- | --- | --- | --- |
| Ala | 3 | 70.1 | A 70%; N 25%; Q 1%; S 1% | Asn | 3 | 84.3 | N 84%; R 1%; S 1% |
| Ser | 6 | 43.3 | S 43%; A 40%; T 12%; I 3% | Thr | 6 | 83.1 | T 83%; M 1%; R 1%; S 1% |
| Ser | 8 | 70.1 | T 70%; S 16%; E 10%; G 1% | Thr | 8 | 77.1 | T 77%; S 7%; N 1%; V 1% |
| Thr | 9 | 79.1 | T 79%; P 7%; V 6%; S 3%; A 1%; G 1% | Val | 9 | 84.3 | V 84%; G 1%; S 1% |
| Cys | 12 | 94.0 | C 94%; V 4% | Val | 12 | 83.1 | V 83%; M 1%; R 1%; S 1%; T 1% |
| Ala | 14 | 86.6 | A 87%; S 12% | Gly | 14 | 59.0 | G 59%; A 20%; S 6%; K 1% |
| Tyr | 15 | 91.0 | Y 91%; H 6%; F 1% | His | 15 | 59.0 | H 59%; Y 22%; A 1%; E 1%; G 1%; N 1%; S 1% |
| Ser | 19 | 73.1 | S 73%; N 19%; T 4%; D 1% | Asp | 19 | 81.9 | N 82%; D 1%; G 1%; K 1%; L 1% |
| Thr | 21 | 68.7 | T 69%; N 16%; S 4%; A 3%; D 3%; H 3% | Ser | 21 | 84.3 | S 84%; A 1%; E 1%; G 1% |
| Thr | 25 | 61.2 | T 61%; S 37% | Ser | 25 | 80.7 | S 81%; G 4%; D 1%; T 1%; V 1% |
| Pro | 26 | 86.6 | P 87%; Q 7%; E 1%; G 1%; L 1% | Leu | 26 | 83.1 | L 83%; V 2%; P 1%; X 1% |
| Asp | 27 | 74.6 | D 75%; E 22%; P 1% | Glu | 27 | 75.9 | E 76%; D 6%; P 2%; Q 2%; T 1% |
| Arg | 31 | 77.6 | R 78%; K 19%; L 3% | Lys | 31 | 78.3 | K 78%; R 6%; A 1%; H 1%; I 1% |
| His | 32 | 53.7 | H 54%; N 37%; S 7%; A 1% | Leu | 32 | 74.7 | L 75%; Q 7%; M 2%; N 2%; R 1% |
| Leu | 33 | 82.1 | L 82%; F 13%; Q 3%; I 1% | Lys | 33 | 81.9 | K 82%; Q 2%; E 1%; L 1%; R 1% |
| Lys | 35 | 95.5 | K 96%; S 3%; R 1% | Arg | 35 | 60.2 | R 60%; K 27%; Q 1% |
| Tyr | 36 | 83.6 | Y 84%; F 15%; H 1% | Trp | 36 | 84.3 | W 84%; F 1%; Y 1% |
| His | 38 | 65.7 | P 66%; H 18%; F 6%; Y 6%; W 3%; L 1% | Ser | 38 | 79.5 | S 80%; A 4%; L 4%; P 1%; T 1% |
| Ser | 48 | 77.6 | S 78%; T 22% | Thr | 48 | 84.3 | T 84%; S 10%; P 1% |
| Trp | 50 | 100 | W 100% | Leu | 50 | 83.1 | L 83%; W 12% |
| Ile | 99 | 98.5 | I 99%; L 1% | Val | 99 | 88.0 | V 88%; I 10% |
| Ala | 132 | 73.1 | A 73%; S 25%; G 1% | Gln | 132 | 96.4 | Q 96%; E 1% |
| Arg | 143 | 97.0 | R 97%; S 3% | Lys | 143 | 72.3 | K 72%; R 25% |
| Val | 153 | 92.5 | V 93%; I 4%; S 1%; W 1% | Ile | 153 | 66.3 | I 66%; V 33% |
| Ile | 165 | 88.1 | I 88%; L 10%; S 1% | Leu | 165 | 71.1 | L 71%; I 25% |
| Leu | 166 | 71.6 | L 72%; I 15%; T 6%; V 6%; S 1% | Thr | 166 | 90.4 | T 90%; S 4%; C 2% |
| Glu | 192 | 55.2 | E 55%; D 45% | Asp | 192 | 98.8 | D 99%; E 1% |
| Leu | 220 | 47.8 | L 48%; I 30%; V 19%; T 3% | Met | 220 | 86.7 | M 87%; I 10%; V 4% |
| Ile | 222 | 79.1 | I 79%; V 21% | Val | 222 | 98.8 | V 99%; I 1% |
| Ala | 224 | 67.2 | A 67%; V 19%; I 13% | Val | 224 | 83.1 | V 83%; I 16%; L 1% |
| Thr | 225 | 62.7 | T 63%; A 36%; S 1% | Ala | 225 | 97.6 | A 98%; G 1%; N 1% |
| Ser | 229 | 50.7 | S 51%; N 30%; G 15%; T 3%; A 1% | Asn | 229 | 97.6 | N 98%; F 1%; G 1% |
| Gln | 238 | 82.1 | Q 82%; E 18% | Glu | 238 | 98.8 | E 99%; N 1% |
| Ala | 240 | 98.5 | A 99%; V 1% | Val | 240 | 89.2 | V 89%; A 10%; K 1% |
| Asp | 245 | 68.7 | D 69%; E 31% | Glu | 245 | 98.8 | E 99%; D 1% |
| Lys | 246 | 100 | K 100% | Arg | 246 | 91.6 | R 92%; K 7%; L 1% |
| Val | 269 | 100 | V 100% | Ile | 269 | 100.0 | I 100% |
| Leu | 273 | 92.5 | L 93%; A 3%; I 3%; M 1% | Ile | 273 | 91.6 | I 92%; M 6%; A 1%; L 1% |
| Lys | 365 | 89.6 | K 90%; R 10% | Arg | 365 | 98.8 | R 99%; K 1% |
| Ile | 369 | 64.2 | I 64%; V 21%; L 12%; Q 1%; Y 1% | Leu | 369 | 89.2 | L 89%; V 8%; I 1% |
| Asp | 373 | 65.7 | D 66%; E 30%; G 1%; H 1%; K 1% | Glu | 373 | 91.6 | E 92%; D 6%; R 1%; S 1% |
| Phe | 376 | 43.3 | I 43%; L 15%; V 10%; F 9%; N 6%; S 6%; H 3%; M 3%; Q 1%; T 1%; Y 1% | Thr | 376 | 56.6 | T 57%; S 34%; N 6%; A 1%; Q 1% |
| Ser | 383 | 74.6 | S 75%; D 13%; T 7%; A 3%; V 1% | Thr | 383 | 91.6 | T 92%; N 4%; S 2%; D 1%; I 1% |
| Glu | 392 | 97.0 | E 97%; G 3% | Ile | 392 | 62.7 | I 63%; M 23%; E 8%; L 2%; V 2%; T 1% |
| Leu | 396 | 74.6 | L 75%; T 10%; M 7%; Q 4%; S 3% | His | 396 | 83.1 | H 83%; Y 5%; L 4%; Q 4%; F 2%; C 1%; T 1% |
| Asn | 398 | 94.0 | N 94%; G 3%; S 3% | Asp | 398 | 97.6 | D 98%; E 1%; N 1% |
| Ile | 402 | 79.1 | V 79%; I 16%; T 4% | Val | 402 | 79.5 | V 80%; I 19% |
| Arg | 403 | 73.1 | R 73%; K 24%; N 3% | Lys | 403 | 88.0 | K 88%; N 6%; R 4%; V 1% |
| Ser | 404 | 47.8 | A 48%; S 19%; P 18%; C 15% | Cys | 404 | 95.2 | C 95%; S 2%; E 1% |
| Gly | 405 | 97.0 | G 97%; S 3% | His | 405 | 69.9 | H 70%; S 24%; T 2%; E 1%; K 1% |
| Phe | 407 | 92.5 | Y 93%; F 6%; H 1% | Tyr | 407 | 94.0 | Y 94%; N 2%; F 1%; H 1% |
| Ser | 424 | 94.0 | S 94%; A 6% | Ala | 424 | 69.9 | A 70%; S 30% |
| Phe | 427 | 79.1 | F 79%; Y 21% | Tyr | 427 | 89.2 | Y 89%; F 11% |
| Thr | 430 | 38.8 | A 39%; T 33%; S 27%; N 1% | Ala | 430 | 92.8 | A 93%; T 4%; V 2%; S 1% |
| Thr | 447 | 77.6 | T 78%; C 22% | Cys | 447 | 98.8 | C 99%; V 1% |
| Asn | 456 | 98.5 | N 99%; S 1% | Asp | 456 | 98.8 | D 99%; E 1% |
| Val | 459 | 100 | V 100% | Leu | 459 | 94.0 | L 94%; V 4%; I 2% |
| Arg | 460 | 91.0 | R 91%; K 9% | Lys | 460 | 96.4 | K 96%; A 2%; R 1% |
| Asn | 461 | 61.2 | S 61%; N 24%; G 15% | Gly | 461 | 95.2 | G 95%; N 2%; S 2% |
| Val | 465 | 97.0 | V 97%; I 1%; P 1% | Ile | 465 | 97.6 | I 98%; S 1%; V 1% |
| Arg | 476 | 68.7 | R 69%; K 31% | Lys | 476 | 94.0 | K 94%; T 2%; H 1%; R 1%; S 1% |
| Ser | 499 | 80.6 | S 81%; T 18%; I 1% | Thr | 499 | 95.2 | T 95%; S 4%; I 1% |
| Ala | 501 | 97.0 | A 97%; S 3% | Asn | 501 | 96.4 | N 96%; K 2%; A 1% |
| Ser | 503 | 52.2 | S 52%; A 21%; G 3%; P 3% | Pro | 503 | 90.4 | P 90%; S 2%; N 1% |
| Ala | 506 | 97.0 | A 97% | Thr | 506 | 92.8 | T 93%; S 4%; R 2%; H 1% |
| Ala | 507 | 83.6 | A 84%; P 6%; S 4%; V 3% | Ser | 507 | 97.6 | S 98%; A 2% |
| Val | 508 | 94.0 | V 94%; L 3% | Met | 508 | 92.8 | M 93%; A 2%; L 2%; H 1%; S 1% |
| Gly | 509 | 100 | G 100% |  |  |  |  |
| Asn | 510 | 94.0 | N 94%; A 3%; S 3% | Ser | 509 | 88.0 | S 88%; A 5%; G 5%; N 1%; P 1% |
| Asn | 526 | 85.1 | N 85%; A 10%; S 3%; T 1% | Thr | 525 | 97.6 | T 98%; A 1%; N 1% |
| Tyr | 527 | 100 | Y 100% | His | 526 | 95.2 | H 95%; Y 4%; F 1% |
| Val | 528 | 100 | V 100% | Ile | 527 | 72.3 | I 72%; V 28% |
| Thr | 532 | 97.0 | T 97%; S 3% | Ser | 531 | 78.3 | S 78%; N 17%; T 2%; G 1%; K 1% |
| Arg | 534 | 100 | R 100% | Lys | 533 | 97.6 | K 98%; R 2% |
| Gly | 539 | 73.1 | G 73%; S 12%; P 10%; E 1%; N 1%; T 1% | Pro | 538 | 55.4 | P 55%; S 31%; A 8%; Q 4%; K 1% |
| Pro | 540 | 74.6 | P 75%; A 12%; S 6%; V 4%; G 1%; N 1% | Gly | 539 | 96.4 | G 96%; A 2%; Q 1% |
| Glu | 543 | 67.2 | E 67%; D 31%; N 1% | Gln | 542 | 90.4 | Q 90%; E 6%; H 2%; D 1% |
| Leu | 546 | 64.2 | M 64%; L 34%; N 1% | Met | 545 | 98.8 | M 99%; L 1% |
| Lys | 550 | 98.5 | K 99%; R 1% | Arg | 549 | 98.8 | R 99%; N 1% |
| Thr | 554 | 100 | T 100% | Ser | 553 | 69.9 | S 70%; T 30% |
| Arg | 556 | 98.5 | R 99%; Q 1% | Ser | 555 | 65.1 | S 65%; R 31%; N 2%; K 1% |
| Arg | 567 | 100 | R 100% | His | 566 | 90.4 | H 90%; R 8%; V 1% |
| Thr | 569 | 88.1 | T 88%; S 6%; A 3%; N 3% | Asn | 568 | 95.2 | N 95%; A 2%; D 1%; S 1% |
| Pro | 571 | 89.6 | P 90%; L 10% | Met | 570 | 62.7 | L 63%; P 31%; M 4%; A 1%; I 1% |
| Lys | 572 | 88.1 | K 88%; R 12% | Arg | 571 | 84.3 | R 84%; K 14%; S 1% |
| Val | 577 | 44.8 | V 45%; I 36%; N 12%; K 3%; F 1%; M 1%; S 1% | Asn | 576 | 43.4 | N 43%; H 42%; V 6%; K 4%; Y 2%; D 1%; Q 1% |
| Asp | 579 | 70.1 | D 70%; E 28%; V 1% | Glu | 578 | 89.2 | E 89%; S 4%; D 2%; V 2%; K 1%; T 1% |
| Ser | 582 | 47.8 | A 48%; T 36%; S 13%; N 3% | Ala | 581 | 78.3 | A 78%; S 14%; T 4%; K 1%; N 1%; V 1% |
| Arg | 583 | 52.2 | K 52%; R 45%; H 3% | Asn | 582 | 90.4 | N 90%; K 5%; R 4%; S 1% |
| Met | 585 | 47.8 | M 48%; L 18%; A 13%; V 10%; I 7%; G 3% | Ile | 584 | 91.6 | I 92%; A 6%; V 2% |
| Glu | 586 | 76.1 | E 76%; D 24% | Lys | 585 | 71.1 | K 71%; N 17%; E 5%; G 2%; T 2%; D 1%; S 1% |
| Asp | 590 | 100 | D 100% | Asn | 589 | 91.6 | N 92%; D 6%; G 2% |
| Val | 596 | 68.7 | V 69%; C 31% | Cys | 595 | 98.8 | C 99%; V 1% |
| Lys | 605 | 100 | K 100% | Ile | 604 | 77.1 | I 77%; T 19%; Q 2%; S 1% |
| Met | 608 | 56.7 | T 57%; M 39%; V 3%; I 1% | Ala | 607 | 98.8 | A 99%; R 1% |
| Gly | 609 | 94.0 | G 94%; S 4%; D 1% | Ser | 608 | 88.0 | S 88%; A 11%; D 1% |
| Ile | 611 | 100 | I 100% | Val | 610 | 67.5 | V 67%; I 33% |
| Gln | 612 | 74.6 | Q 75%; E 16%; K 7%; R 1% | Lys | 611 | 97.6 | K 98%; M 2% |
| Asp | 616 | 80.6 | D 81%; A 12%; E 7% | Gln | 615 | 92.8 | Q 93%; L 5%; E 1%; H 1% |
| Ile | 633 | 89.6 | I 90%; V 10% | Val | 632 | 97.6 | V 98%; E 1% |
| Glu | 644 | 97.0 | E 97%; D 3% | Gln | 643 | 59.0 | Q 59%; E 29%; L 4%; S 2%; D 1%; H 1%; N 1% |
| Asn | 645 | 59.7 | N 60%; D 15%; S 15%; E 10% | Asp | 644 | 81.9 | D 82%; E 14%; N 1% |
| Glu | 647 | 64.2 | E 64%; D 31%; Q 4% | Asp | 646 | 94.0 | D 94%; E 2%; S 1% |
| Ala | 649 | 44.8 | A 45%; T 40%; S 12%; V 3% | Thr | 648 | 59.0 | T 59%; S 28%; D 5%; A 4%; H 1%; N 1%; V 1% |
| Asp | 650 | 65.7 | D 66%; G 24%; R 4%; E 1%; N 1%; S 1%; T 1% | Ser | 649 | 56.6 | S 57%; T 17%; A 11%; K 5%; F 2%; C 1%; D 1%; E 1%; G 1%; L 1%; R 1% |
| Arg | 651 | 88.1 | R 88%; K 6%; C 3%; L 3% | Lys | 650 | 88.0 | K 88%; M 6%; L 2%; R 2% |
| Tyr | 653 | 83.6 | Y 84%; F 16% | Phe | 652 | 97.6 | F 98%; W 1% |
| Asp | 660 | 100 | D 100% | Glu | 659 | 95.2 | E 95%; D 4%; G 1% |
| Pro | 662 | 97.0 | P 97%; S 3% | Asn | 661 | 69.9 | S 70%; N 27%; A 2%; G 1% |
| Leu | 663 | 79.1 | L 79%; P 16%; S 3%; M 1% | Pro | 662 | 81.9 | P 82%; L 16%; E 1%; V 1% |
| Ala | 664 | 61.2 | A 61%; P 19%; T 6%; G 3%; H 3%; Y 3%; Q 1%; S 1%; V 1% | Ser | 663 | 63.9 | S 64%; A 23%; P 6%; Q 5%; H 1%; V 1% |
| Glu | 665 | 92.5 | E 93%; Q 6%; D 1% | Ala | 664 | 96.4 | A 96%; E 2%; F 1% |
| Arg | 671 | 89.6 | R 90%; K 10% | Leu | 670 | 74.7 | L 75%; H 14%; T 6%; M 2%; R 1%; W 1% |
| Arg | 672 | 86.6 | R 87%; K 10%; H 3% | Asn | 671 | 78.3 | N 78%; H 18%; D 1%; T 1%; V 1% |
| Cys | 674 | 89.6 | C 90%; S 9%; R 1% | Arg | 673 | 98.8 | R 99%; S 1% |
| Tyr | 690 | 65.7 | Y 66%; F 34% | Phe | 689 | 95.2 | F 95%; Y 5% |
| Tyr | 694 | 49.3 | F 49%; Y 46%; C 3%; N 1% | Phe | 693 | 94.0 | F 94%; M 2%; D 1%; L 1%; S 1% |
| Ala | 714 | 100 | A 100% | Ser | 713 | 61.4 | A 61%; S 39% |
| Leu | 783 | 98.5 | L 99% | Phe | 782 | 97.6 | F 98%; M 2% |
| Asp | 818 | 83.6 | D 84%; G 13%; N 1% | Asn | 817 | 92.8 | N 93%; D 2%; E 2%; K 1%; S 1% |
| Arg | 819 | 76.1 | R 76%; K 21% | Lys | 818 | 98.8 | K 99% |
| Ser | 823 | 91.0 | S 91%; T 7% | Asn | 822 | 94.0 | N 94%; S 4%; R 1% |
| Met | 838 | 73.1 | M 73%; L 27% | Leu | 837 | 98.8 | L 99%; V 1% |
| Gly | 842 | 95.5 | G 96%; A 3% | Cys | 841 | 96.4 | C 96%; G 2%; V 1% |
| Met | 857 | 65.7 | L 66%; M 27%; I 6% | Ile | 856 | 91.6 | I 92%; V 4%; T 2%; M 1% |
| Tyr | 858 | 97.0 | Y 97%; F 1% | Ala | 857 | 94.0 | A 94%; L 2%; F 1%; V 1% |
| Glu | 860 | 56.7 | E 57%; D 39%; P 1%; S 1% | Asp | 859 | 85.5 | D 86%; E 13% |
| Asp | 861 | 86.6 | D 87%; E 9%; T 3% | Gly | 860 | 86.7 | G 87%; D 11%; E 1% |
| Gly | 864 | 62.7 | H 63%; G 12%; M 9%; E 4%; N 4%; S 4%; A 1%; Q 1% | Arg | 863 | 88.0 | R 88%; K 4%; M 4%; Q 4% |
| Tyr | 867 | 79.1 | Y 79%; F 19%; L 1% | Phe | 866 | 90.4 | F 90%; L 7%; Y 2% |
| His | 868 | 49.3 | S 49%; Y 30%; H 13%; N 6%; Q 1% | Tyr | 867 | 98.8 | Y 99%; S 1% |
| Thr | 871 | 70.1 | T 70%; S 30% | Ser | 870 | 100.0 | S 100% |
| Met | 874 | 100 | M 100% | Leu | 873 | 98.8 | L 99%; M 1% |
| Thr | 877 | 71.6 | T 72%; S 10%; N 9%; H 4%; G 3%; A 1% | Lys | 876 | 90.4 | K 90%; S 5%; A 2%; R 2% |
| His | 880 | 91.0 | N 91%; H 4%; S 4% | Asn | 879 | 98.8 | N 99%; S 1% |
| His | 882 | 58.2 | D 58%; H 22%; E 16%; N 3% | Asp | 881 | 94.0 | D 94%; E 5%; H 1% |
| Leu | 886 | 43.3 | V 43%; L 31%; I 18%; H 4%; M 3% | Val | 885 | 91.6 | V 92%; L 7%; I 1% |
| Glu | 889 | 97.0 | E 97%; D 3% | Ala | 888 | 66.3 | A 66%; V 16%; H 6%; E 5%; L 5%; G 1%; N 1% |
| Ile | 890 | 61.2 | V 61%; I 39% | Val | 889 | 57.8 | I 58%; V 42% |
| Ala | 893 | 76.1 | A 76%; S 24% | Ser | 892 | 98.8 | S 99%; D 1% |
| Glu | 895 | 61.2 | E 61%; V 16%; P 12%; Q 7%; F 1%; T 1% | Tyr | 894 | 98.8 | Y 99%; H 1% |
| Met | 923 | 49.3 | L 49%; V 22%; M 16%; I 12% | Leu | 922 | 80.7 | L 81%; M 18%; V 1% |
| Val | 929 | 83.6 | V 84%; S 16% | Glu | 928 | 98.8 | E 99%; K 1% |
| Leu | 934 | 67.2 | L 67%; V 21%; M 9%; A 3% | Val | 933 | 84.3 | V 84%; L 14%; M 1% |
| Asp | 951 | 88.1 | D 88%; E 12% | Glu | 950 | 98.8 | E 99%; P 1% |
| Met | 955 | 100 | M 100% | Leu | 954 | 77.1 | L 77%; I 16%; V 6%; P 1% |
| Lys | 958 | 100 | K 100% | Gln | 957 | 98.8 | Q 99%; R 1% |
| Leu | 959 | 100 | L 100% | Ile | 958 | 98.8 | I 99%; V 1% |
| Lys | 960 | 41.8 | R 42%; T 31%; Q 22%; K 3%; E 1% | Thr | 959 | 100.0 | T 100% |
| Ala | 961 | 67.2 | A 67%; H 16%; P 16% | Pro | 960 | 100.0 | P 100% |
| Asp | 963 | 73.1 | D 73%; N 21%; T 4%; S 1% | Asn | 962 | 95.2 | N 95%; D 5% |
| Gly | 979 | 65.7 | G 66%; L 33%; A 1% | Leu | 978 | 98.8 | L 99%; F 1% |
| Leu | 980 | 83.6 | L 84%; I 16% | Met | 979 | 65.1 | M 65%; L 34%; I 1% |
| Ile | 983 | 44.8 | I 45%; L 33%; V 18%; A 4% | Thr | 982 | 92.8 | T 93%; L 6%; I 1% |
| Ile | 987 | 49.3 | V 49%; I 48%; A 1%; F 1% | Val | 986 | 92.8 | V 93%; A 4%; F 2%; I 1% |
| Gly | 994 | 53.7 | G 54%; D 22%; A 9%; E 3%; Q 3%; V 3%; C 1%; P 1%; T 1% | Pro | 993 | 95.2 | P 95%; F 2%; Q 1%; S 1% |
|  |  |  |  | Ala | 994 | 71.1 | G 71%; A 24%; V 2%; E 1%; S 1% |

**Supplementary Table 2.**
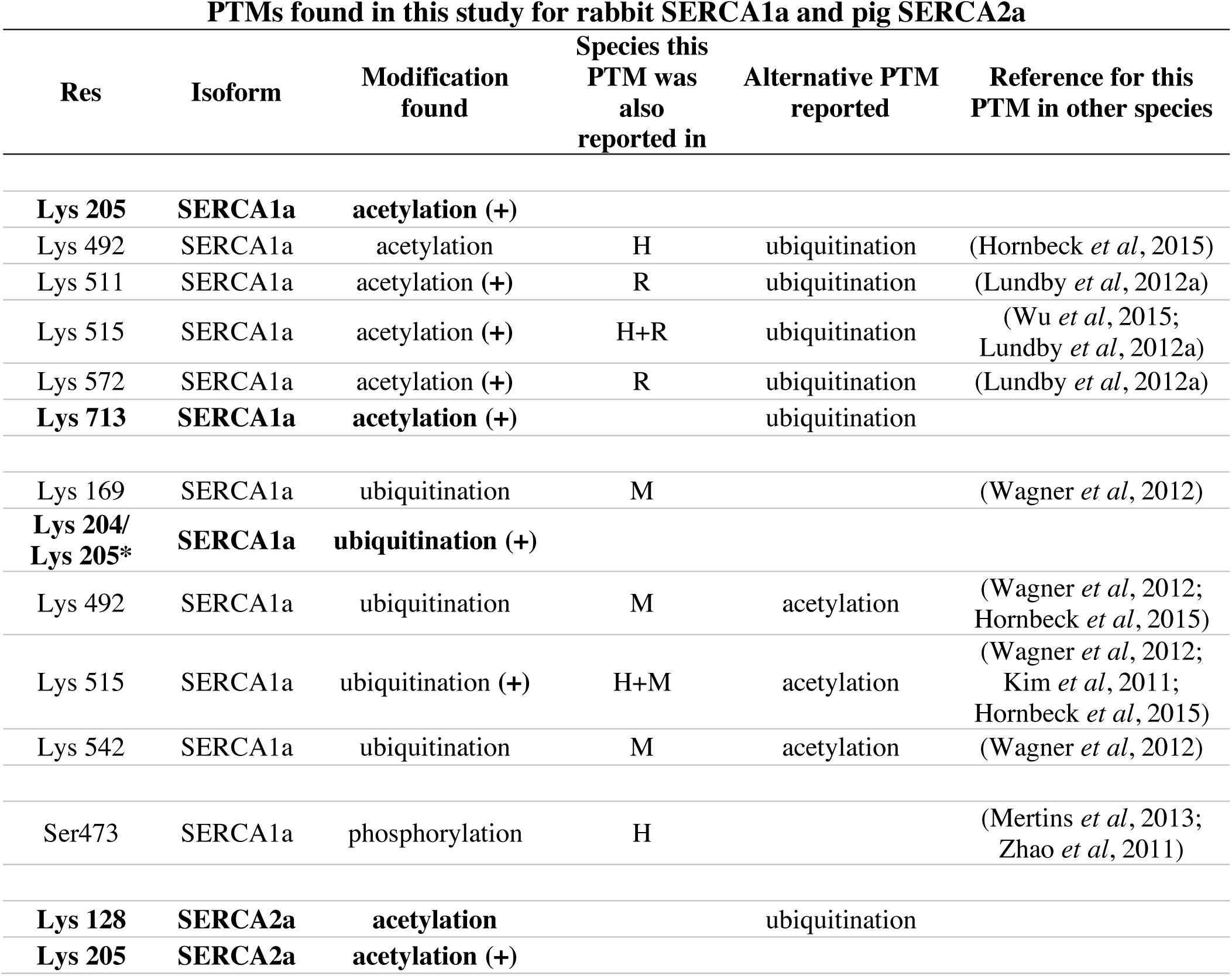
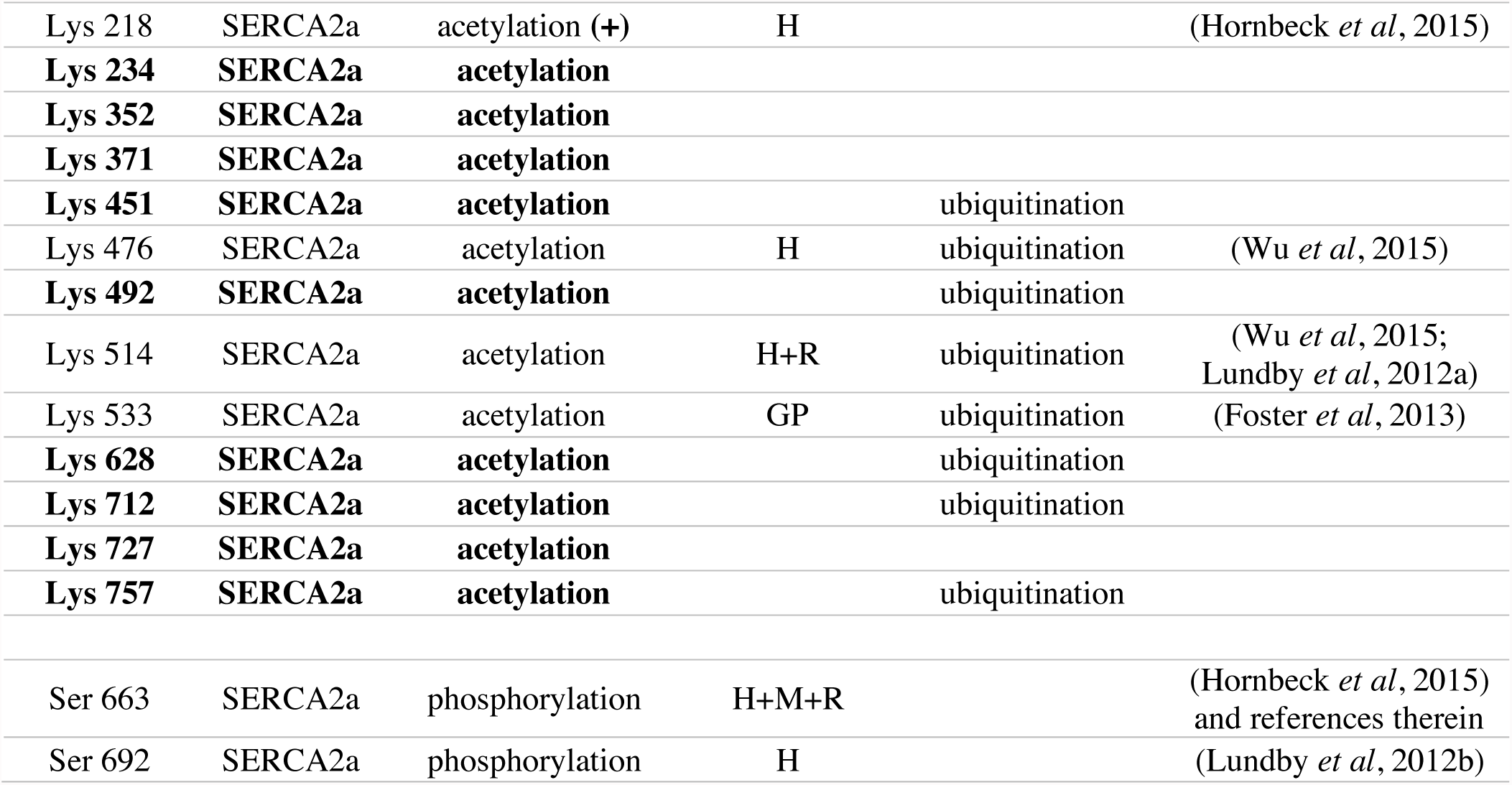
A list of PTMs found in the MS analysis of purified SERCA1a and SERCA2a. Only peptides expressing a high identification confidence and a localization confidence of 99%+ were chosen and are shown in this table. PTMs that have not been previously reported elsewhere are highlighted in bold. PTMs that were found alternatively modified either in this study or elsewhere are indicated with (+). The species where the previously reported PTMs were identified are also indicated (H – humans, R – rats, M – mice, GP – guinea pig). In SERCA1a either Lys 204 or Lys 205 is ubiquitinated (*).

| PTMs found in this study for rabbit SERCA1a and pig SERCA2a |  |  |  |  |  |
| --- | --- | --- | --- | --- | --- |
| Res | Isoform | Modification found | Species this PTM was also reported in | Alternative PTM reported | Reference for this PTM in other species |
| **Lys 205** | **SERCA1a** | **acetylation (+)** |  |  |  |
| Lys 492 | SERCA1a | acetylation | H | ubiquitination | (Hornbeck *et al*, 2015) |
| Lys 511 | SERCA1a | acetylation (+) | R | ubiquitination | (Lundby *et al*, 2012a) |
| Lys 515 | SERCA1a | acetylation (+) | H+R | ubiquitination | (Wu *et al*, 2015; Lundby *et al*, 2012a) |
| Lys 572 | SERCA1a | acetylation (+) | R | ubiquitination | (Lundby *et al*, 2012a) |
| **Lys 713** | **SERCA1a** | **acetylation (+)** |  | ubiquitination |  |
| Lys 169 | SERCA1a | ubiquitination | M |  | (Wagner *et al*, 2012) |
| **Lys 204/**<br>**Lys 205\*** | **SERCA1a** | **ubiquitination (+)** |  |  |  |
| Lys 492 | SERCA1a | ubiquitination | M | acetylation | (Wagner *et al*, 2012; Hornbeck *et al*, 2015) |
| Lys 515 | SERCA1a | ubiquitination (+) | H+M | acetylation | (Wagner *et al*, 2012; Kim *et al*, 2011; Hornbeck *et al*, 2015) |
| Lys 542 | SERCA1a | ubiquitination | M | acetylation | (Wagner *et al*, 2012) |
| Ser473 | SERCA1a | phosphorylation | H |  | (Mertins *et al*, 2013; Zhao *et al*, 2011) |
| **Lys 128** | **SERCA2a** | **acetylation** |  | ubiquitination |  |
| **Lys 205** | **SERCA2a** | **acetylation (+)** |  |  |  |
| Lys 218 | SERCA2a | acetylation (+) | H |  | (Hornbeck *et al*, 2015) |
| **Lys 234** | **SERCA2a** | **acetylation** |  |  |  |
| **Lys 352** | **SERCA2a** | **acetylation** |  |  |  |
| **Lys 371** | **SERCA2a** | **acetylation** |  |  |  |
| **Lys 451** | **SERCA2a** | **acetylation** |  | ubiquitination |  |
| Lys 476 | SERCA2a | acetylation | H | ubiquitination | (Wu *et al*, 2015) |
| **Lys 492** | **SERCA2a** | **acetylation** |  | ubiquitination |  |
| Lys 514 | SERCA2a | acetylation | H+R | ubiquitination | (Wu *et al*, 2015;<br>Lundby *et al*, 2012a) |
| Lys 533 | SERCA2a | acetylation | GP | ubiquitination | (Foster *et al*, 2013) |
| **Lys 628** | **SERCA2a** | **acetylation** |  | ubiquitination |  |
| **Lys 712** | **SERCA2a** | **acetylation** |  | ubiquitination |  |
| **Lys 727** | **SERCA2a** | **acetylation** |  |  |  |
| **Lys 757** | **SERCA2a** | **acetylation** |  | ubiquitination |  |
| Ser 663 | SERCA2a | phosphorylation | H+M+R |  | (Hornbeck *et al*, 2015)<br>and references therein |
| Ser 692 | SERCA2a | phosphorylation | H |  | (Lundby *et al*, 2012b) |

**Supplementary Table 3.**
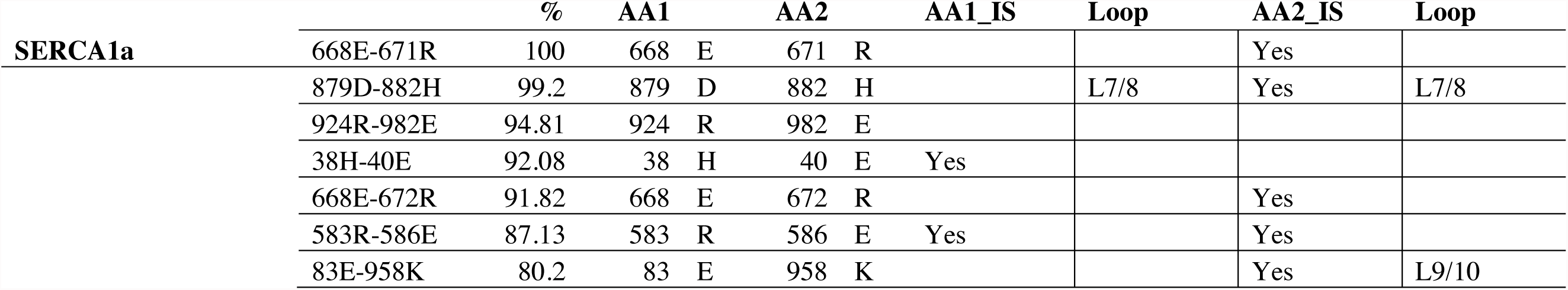
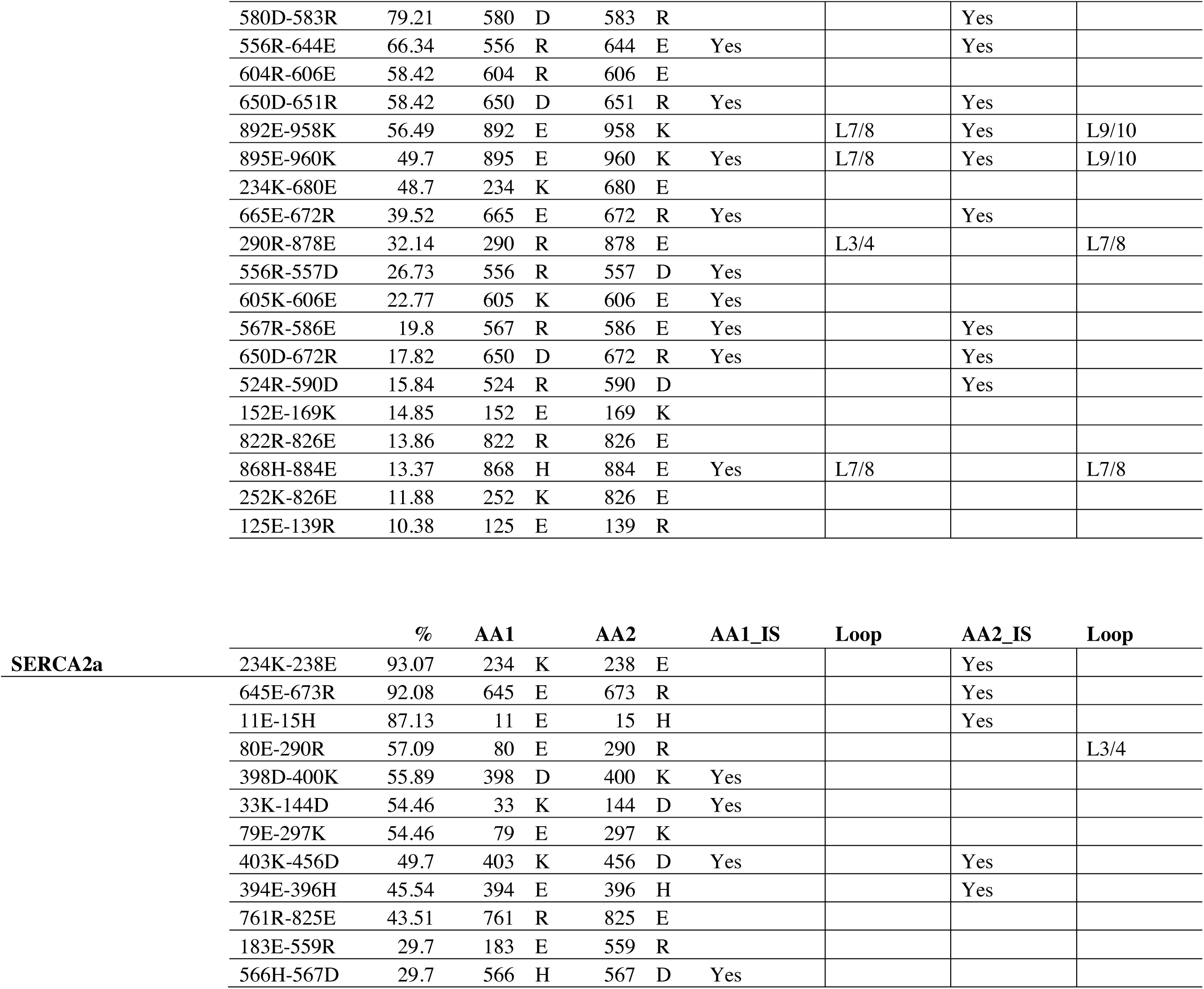
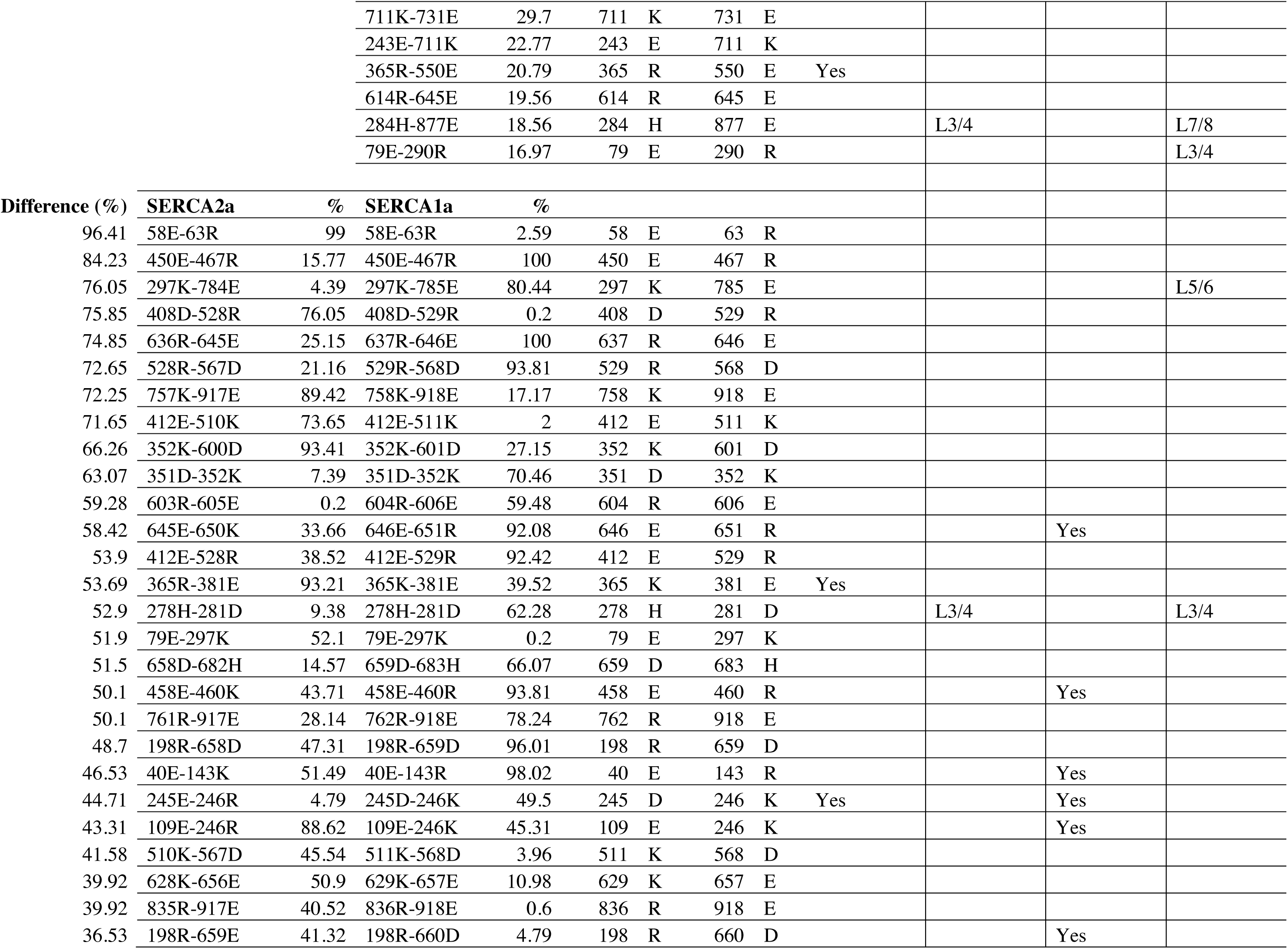
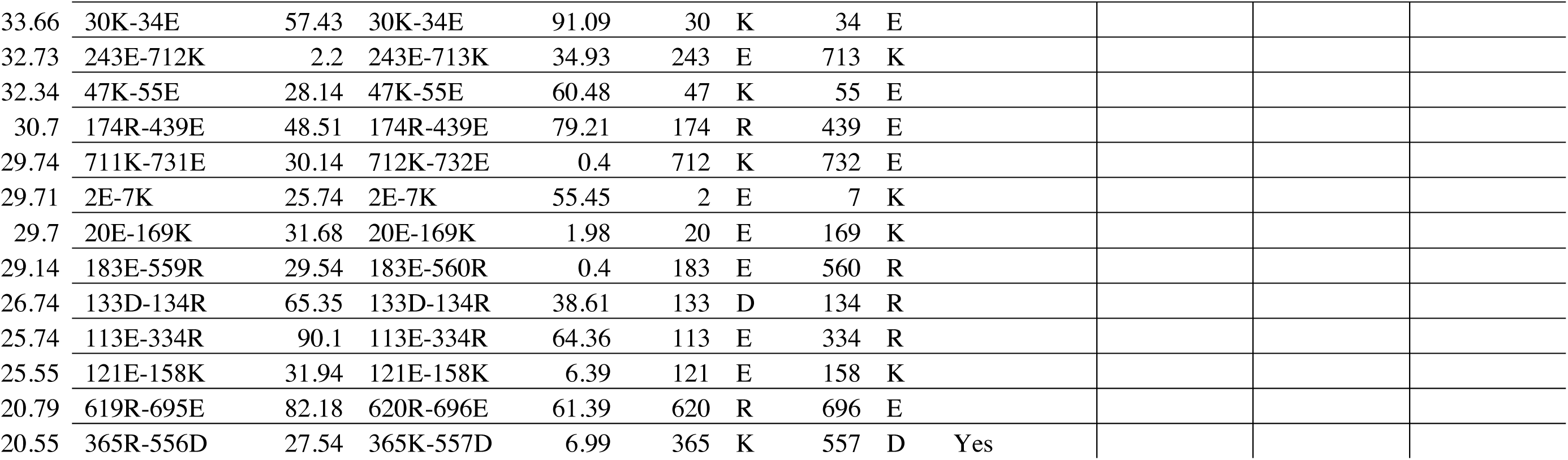
Salt bridges found during MD simulations. Only unique interactions of SERCA1a and SERCA2a present during at least 10% of the total simulation time are shown. Residue pairs that form salt bridges in both isoforms but have a different interaction time of at least 20% are also shown below. AA1 and AA2 stand for the first and second amino acids in the interacting residue pair, IS – isoform specific, % - percent of total simulation time spent interacting.

Author Contributions
Conceptualization, C.O., P.V., P.N.; Methodology, A.S., N.D., I.V., M.D.M., E.W., C.O., P.V., P.N.; Formal analysis, A.S., N.D., J.L.A., R.D., S.S.; Investigation, A.S., N.D., R.D., S.S., J.D.R., I.V., J.C.; Writing – original draft preparation, A.S., P.N., P.V.; Writing – review and editing, A.S., J.L.A., P.N., P.V.; Visualization, A.S. Supervision, C.O., P.V., P.N.; Funding acquisition, A.S., C.O., P.V., P.N.

## Abbreviations

SERCA: Sarcoplasmic reticulum Ca^2+^-ATPase
CPA: Cyclopiazonic acid
C_12_E_8_: octaethylene glycol monododecyl ether

